# Ageing impairs the neuro-vascular interface in the heart

**DOI:** 10.1101/2022.07.29.501999

**Authors:** Julian U. G. Wagner, Lukas Tombor, Pedro Felipe Malacarne, Lisa-Maria Kettenhausen, Josefine Panthel, Maria Cipca, Kathrin A. Stilz, Ariane Fischer, Marion Muhly-Reinholz, Wesley T. Abplanalp, David John, Giulia K. Buchmann, Stephan Angendohr, Ehsan Amin, Katharina Scherschel, Nikolaj Klöcker, Malte Kelm, Dominik Schüttler, Sebastian Clauss, Stefan Guenther, Thomas Boettger, Thomas Braun, Christian Bär, Eleonora Nardini, Selma Osmanagic-Myers, Christian Meyer, Andreas M. Zeiher, Ralf P. Brandes, Guillermo Luxán, Stefanie Dimmeler

## Abstract

Aging is a major risk factor for impaired cardiovascular health. The aging myocardium is characterized by electrophysiological dysfunctions such as a reduced heart rate variability. These alterations can be intrinsic within cardiomyocytes, but might be modulated by the cardiac autonomic nervous system, as well^1^. It is known that nerves align with vessels during development^2^, but the impact of aging on the cardiac neuro-vascular interface is unknown. Here, we report that aging reduces nerve density specifically in the left ventricle and dysregulates vascular-derived neuro-regulatory genes. Aging leads further to a down-regulation of miR-145 and de-repression of the neuro-repulsive factor Semaphorin-3A. miR-145 deletion increased *Sema3a* expression and reduced axon density, thus mimicking the observed aged heart phenotype. Removal of senescent cells, which accumulated with chronological age while nerve density declined, rescued from age-induced dennervation, reduced *Sema3a* expression and preserved heart rate variability. These data suggest that senescence-associated regulation of neuro-regulatory genes contributes to a declined nerve density of the aging heart and thereby to a reduced heart rate variability.

## Main Text

The vasculature and nervous system form complex, highly branched networks, which are frequently interdependent and functionally linked. Vessel-nerve alignments are mediated by nerve-derived signals that act on endothelial cells or, conversely, the formation of nerve fibers along a preformed vessel template^2^. Thereby, guidance cues such as Semaphorins, Eph/ephrins and vascular endothelial growth factor (VEGF)/VEGFR regulate vessels and neurons, and extend or repel axonal growth^3^. Afferent and efferent cardiac neurotransmission via sympathetic and parasympathetic cardiac nerves modulates many physiological functions of the heart. Hence imbalances of either branches can lead to arrhythmias. For instance, impaired cardiac parasympathetic activity is a negative prognostic indicator and can lead to ventricular arrhythmia^4,5^, whereas both excessive^6–8^ and reduced^9^ sympathetic activity can lead to arrhythmias. Moreover, cardiac denervation lead to silent ischemia and lethal arrhythmia in diabetic hearts ^5,9,10^. A reduced heart rate variability is indicative of an impairment of both parasympathetic and sympathetic innervation in the elderly and has a negative prognostic value^11^. Beyond the control of electrical stability, innervation has additional functions and, for example, is essential for regeneration of the heart, as shown in postnatal post-infarction regeneration models in mice^12,13^. In the vasculature and in other organs innervation can control inflammation^14,15^. However, whether aging has an impact on cardiac innervation on a cellular and mechanistic level is unknown.

Here, we explored the impact of aging on nerve density in old mice. We used 18-20 month old male C57Bl/6J mice, which revealed diastolic dysfunction, while ejection fraction was preserved (**Suppl. Fig. 1a, b**). Pan-neuronal staining for Tuj1 showed a robust reduction of axon density in 18-month old mouse hearts compared to 12 week old young mice (**Fig.1a**). The age-induced reduction in nerve density was specifically detected in the left ventricle, while the right ventricular innervation was comparable between old and young mice (**Fig.1b, c**). The extend of age-dependent decline in the epicardial region was not as strong as in the subendocardial and myocardial regions, but high magnification images proofed a robust decline of sub-epicardial axon density in aged hearts as well (**Fig.1b; Suppl. Fig. 2a**). This observation was further confirmed in whole mount staining of old mouse hearts, which showed a decline in nervous fibers across the left posterior wall (**Fig.1d**). To assess the specific time-point when denervation starts, we performed a time course study showing a decline of nerve density already at 16 month of age with a further decline at 22 months (**Fig.1e; Suppl. Fig. 2b**).

**Fig. 1.**
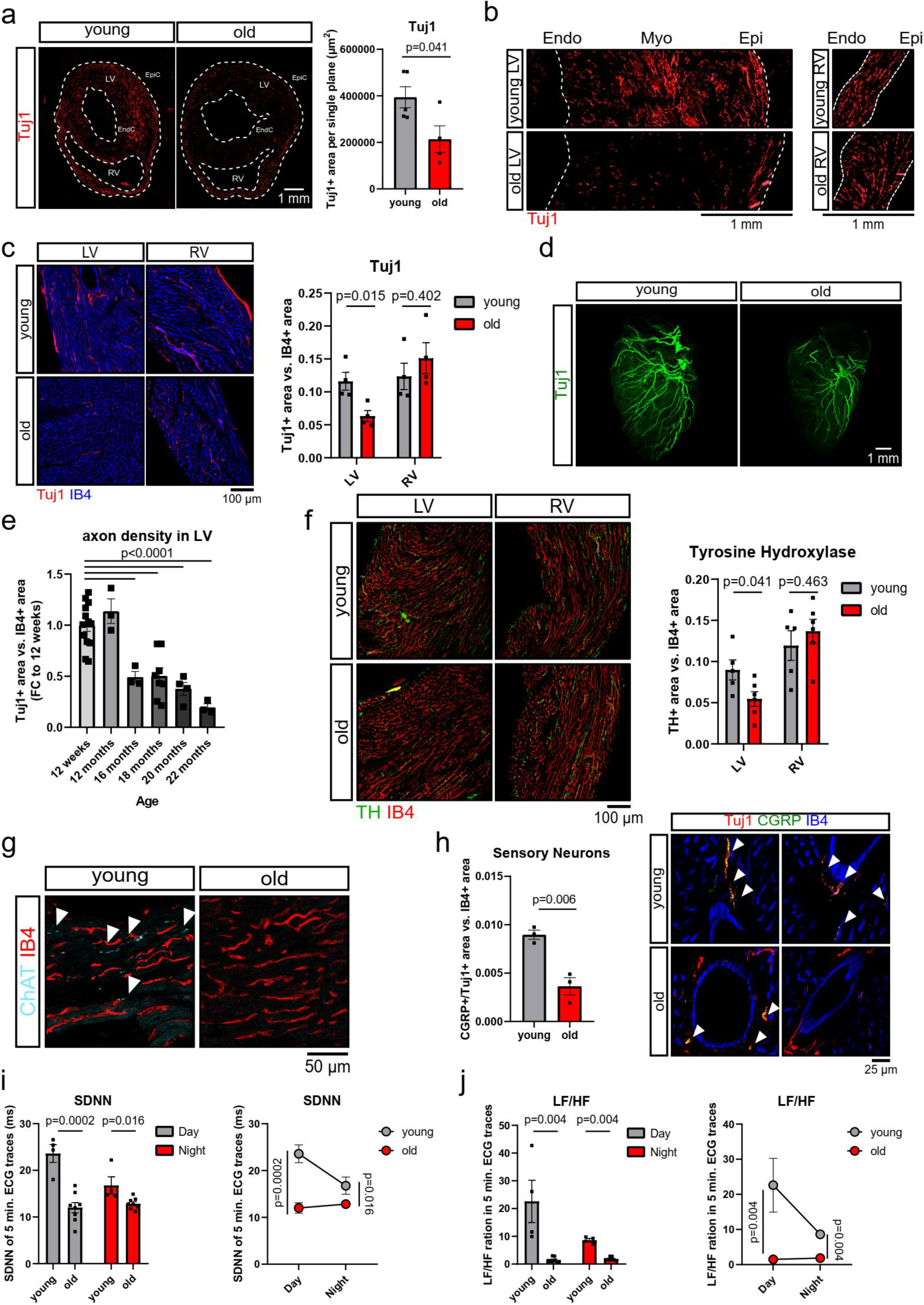
Aging impairs cardiac innervation. **a** Tile scan of male 12-weeks young vs. 18-months old mouse heart cross-sections (LV: left ventricle; RV: right ventricle). Autonomic nervous system is shown by Tuj1 staining (red) and was quantified by Tuj1-positive area per single Z-plane (young: n=5 vs. old: n=4). **b** Zoom-in of the left and right ventricular wall shown in panel a. **c** Quantification of Tuj1-positive area vs. IB4-positive area in young (12 weeks) vs. old (20 months) left (LV) and right ventricles (RV). Innervation was assessed by Tuj1 (red) and normalized to IB4 (blue) (n=4). **d** Whole mount Tuj1 (green) staining of male young (12 weeks) and old (18 months) mouse hearts. One representative image per group is shown (n=3). **e** Left-ventricular innervation (Tuj1=green) vs. IB4 (red) at 12 weeks, 12 months, 16 months, 18 months, 20 months and 22 months of age (n=12 vs. n=3, n=3, n=8, n=4, n=3). **f** Tyrosine hydroxylase (TH) staining (green) vs. IB4 (red) in male young (12 weeks) vs. old (18 and 20 months) left and right ventricles (young n=5 vs. old n=6). **g** Representative choline acetyltransferase (ChAT=cyano) staining in young (12 weeks) vs. old (18 months) old mouse hearts (cardiac base). IB4 (red) served as counter stain (n=3). **h** Immunofluorescence staining of sensory nerves (CGRP=green and Tuj1=red, indicated by white arrow heads) vs. IB4 (blue) in young (12 weeks) vs. old (18 and 20 months) mouse hearts (n=3). **i, j** Heart rate variability was assessed by telemetric ECG recordings in young (15 weeks) vs. old (19 months) mice. Heart rate variability was determined as standard deviation of normal RR-Intervals (SDNN; I) and low frequency to high frequency ratio (LF/HF; j) in ECG traces of 5 minutes at day and night (n=4 vs. n=8). Data are shown as mean and error bars indicate the standard error of the mean (SEM). After passing normality test, statistical power of young vs. old mice was assessed using the unpaired, two-tailed t-test (a, c, f, h, i, j). To compare multiple groups, an ordinary one-way ANOVA test with a post-hoc Tukey test was used (e).

Next, we assessed which types of nerves are affected by aging. The heart is innervated by sympathetic, parasympathetic and sensory fibers, which are commonly stained for tyrosine hydroxylase (TH)^16,17^, choline acetyltransferase (ChAT)^18,19^ and calcitonin gene-related peptide (CGRP)^20^, respectively. TH-positive nerves were present in both ventricles, but were selectively reduced in the left ventricle of aged mice (**Fig.1f**). ChAT-positive nerves were sparse in either ventricles and only occasionally detected in the cardiac base of young hearts (**Fig.1g**). Sensory neurons were exclusively detected in perivascular regions and were also significantly diminished by age (**Fig.1h**). Aging additionally resulted in a higher incidence of ventricular tachycardia and arrhythmias in Langendorff perfused hearts of 18 month old mice compared to young mice (**Suppl. Fig. 3)** documenting increased electrical instability. Together these data demonstrate a decline of cardiac innervation in the left ventricle of old aged mice.

To shed light on the functional consequences that arise from the age-associated cardiac denervation, we assessed heart rate variability by time domain and frequency domain analyses in awake mice using telemetric ECG tracing. In line with a recent report^21^, we observed a reduced variation of the RR-intervals (SDNN) in aged versus young mice (**Fig. 1i**). Especially, frequency-domain analyses as assessed by LF/HF ratio, which can be considered as an indicator for the sympatho-vagal balance^22^, was reduced with age, suggesting a reduced sympathetic activity in aged animals (**Fig. 1j**). The day-night-rhythm was also impaired with age (**Fig. 1i, j**). Taken together, aging leads to left ventricle-specific decrease in neuronal density that correspond to decreased heart rate variability and arrhythmias.

The vasculature and the nervous system co-develop and remain aligned also in the mature heart (**Fig. 2a**). To address if the decline in nerve density may be secondary to age-related capillary rarefaction, we histologically assessed capillary density over 22 months (**Suppl. Fig. 4**). However, capillary density only decreased at 22 months (**Suppl. Fig. 4**). At 16 months, when the initial decline of nerve density was observed, no difference in capillary density was detectable excluding that the reduction of nerves is secondary to the loss of capillaries. However, vascular alignment of nerves is lost in 18-month old mouse hearts (**Fig. 2b, c**), which might have been caused by a dysregulation of neuro-guidance cues in the vasculature.

**Fig. 2.**
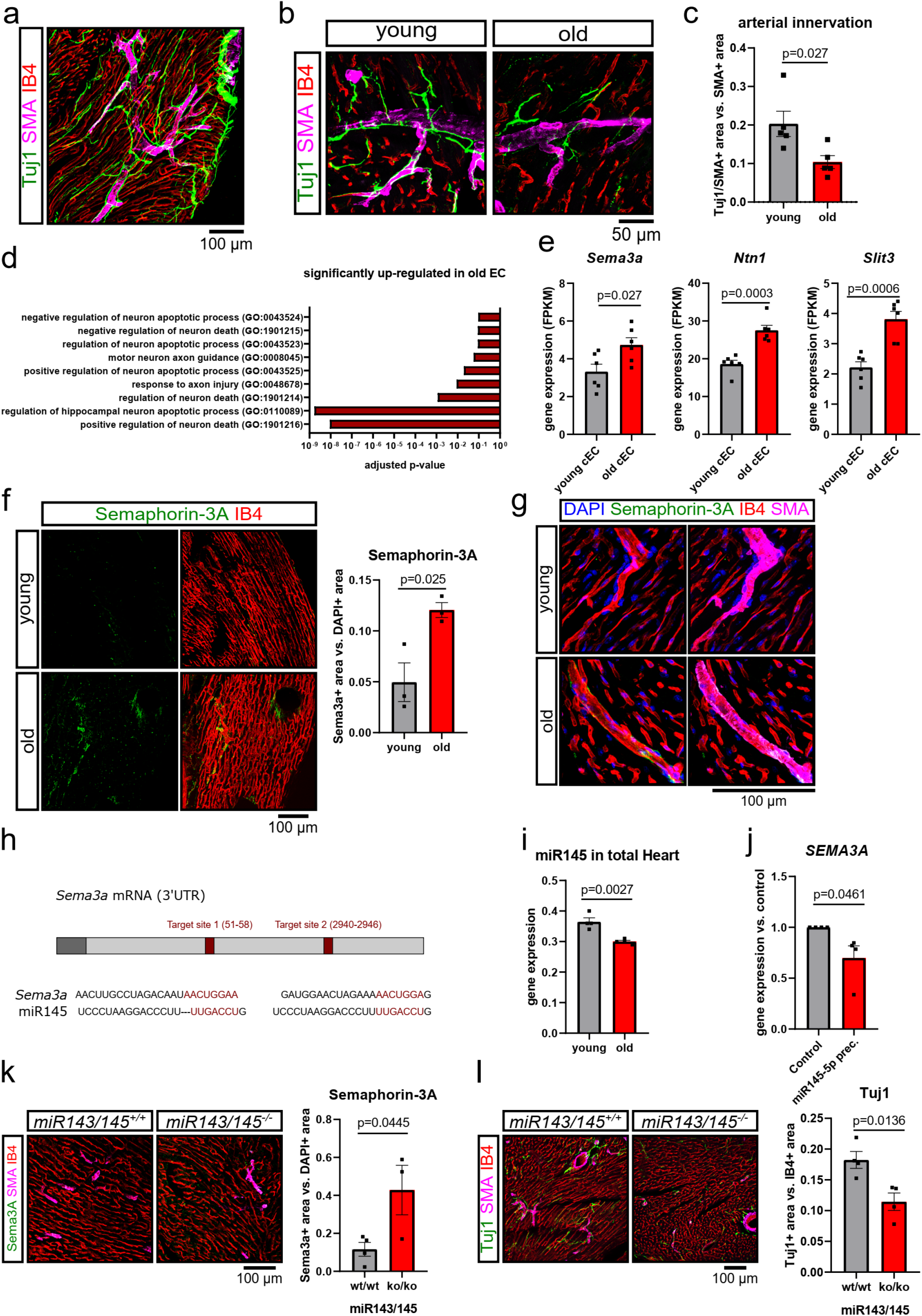
Aging impairs the cardiac neurovascular interface. **a** Representative image of the neurovascular alignment in a young (12 weeks) male mouse heart. Nervous fibers are assessed by Tuj1 staining (green), endothelial cells by IB4 (red) and arterioles by smooth muscle actinin (SMA,magenta) staining. **b, c** Neurovascular alignment in male young (12 weeks) vs. old (18 months) left ventricles as assessed by Tuj1 (green) and SMA (magenta) double positive areas. IB4 (red) served as counter stain (n=5). **d** Gene ontology (GO) analysis of nervous system related GO terms in bulk RNA sequencing data of isolated cardiac endothelial cells from young (12 weeks) vs. old (20 months) mice (n=6). GO term analysis were performed using the online platform *Enrichr*. **e** Gene expression of *Sema3a, Ntn1* and *Slit3* mRNA (FPKM) in young (12 weeks) vs. old (20 months) isolated cardiac mouse endothelial cells (n=6). **f, g** Immunofluorescence staining of Semaphorin-3A (green) and IB4 (red) male in young (12 weeks) and old (20 months) left ventricles. An overview image including the quantification is shown in panel f while panel g shows representative high magnification images (n=3). **h** Schematic representation of miR145 binding-sites described in the *Sema3a* 3’UTR^26^. **i** Expression of miR-145 in total hearts from young and old mice as assessed via microarray (n=4). **j** Relative *Sema3a* mRNA expression in human umbilical cord vein endothelial cells (HUVEC) upon miR145-5p precursor transfection (n=4). **k** Immunofluorescence staining of Semaphorin-3A (green) in hearts derived from young (13-14 weeks) male and female miR143/145-knockout mice. SMA (magenta) and IB4 (red) served as control staining (n=4). **l** Immunofluorescence staining of Tuj1 (green) in hearts derived from young (13-14 weeks) male and female miR143/145-knockout mice. SMA (magenta) and IB4 (red) served as control staining (n=4). Data are shown as mean and error bars indicate the standard error of the mean (SEM). After passing normality test, statistical power was assessed using the unpaired, two-tailed t-test.

To address, if neuro-guidance cues might be dysregulated in the cardiac endothelium of the aging heart, we isolated endothelial cells from young and old mice and performed RNA sequencing. GO term analysis of significantly induced genes demonstrated pathways assigned to “neuronal death” and “axon injury”, (**Fig. 2d**). Genes in these pathways include Semaphorin-3a (*Sema3a)*, which patterns the autonomic nervous system during development^23^. Semaphorin-3A is further essential to maintain normal heart rhythm through sympathetic innervation patterning, but induces vulnerability to arrhythmias if overexpressed in cardiomyocytes^6^ (**Fig. 2e**). In addition, we found upregulation of members of the Slit/Robo family, such as *Slit3*, which can mediate repulsive signals^24^, and *Netrin-1 (Ntn1*), a laminin-related secreted protein, which may switch attraction to repulsive responses in a dose-dependent manner^25^ (**Fig. 2e**). Interestingly, recent studies suggest that the combination of guidance factors (such as Slit-family members and Netrin-1) can act in concert to modulate cellular responses^25^. Validation of protein expression confirmed the upregulation of Semaphorin-3A in aged mouse hearts (**Fig. 2f**) and further showed that Semaphorin-3A is predominantly expressed by vascular cells (**Fig. 2g**).

Since both overexpression and deletion of *Sema3a* may lead to sudden cardiac death and ventricular fibrillation^7^, we investigated up-stream pathways, which may control age-induced induction of *Sema3a. Sema3a*-mRNA has two miR-145 binding sites in the 3’UTR^26^ (**Fig. 2h**), and miR-145-5p was significantly reduced in the aging heart (**Fig. 2i**). Therefore, we hypothesize that miR-145 might repress Sema3a in the young heart. Overexpression of miR-145 indeed repressed *Sema3a* in human umbilical cord vein endothelial cells *in vitro* (**Fig. 2j**). Furthermore, *miR-143/145*^*-/-*^ mice showed increased levels of Semaphorin-3a among vessels (**Fig. 2k**) and reduced axon density (**Fig. 2l**) even at young age (10-15 weeks). Together, these data suggest that loss of miR-145 induced de-repression of *Sema3a* is sufficient to reduce cardiac nerve density.

Importantly, *SEMA3A* was also up-regulated in senescent endothelial cells, which were generated by continuous passaging to induce replicative senescence as evidenced by acidic β-galactosidase staining (**Fig. 3a, b**). Interestingly, cellular senescence is induced concomitantly in the aging mouse hearts when neuronal density declines at 16 months (**Fig. 3c-e**). Moreover, genetic models of premature senescence such as 4^th^ generation *Tert*^*-/-*^ mice that lack telomerase confirmed a decline in nerve density (**Fig.3f, g**). By applying a senescent score^27^ to our previously published single nuclei RNA sequencing data of young vs. old mouse hearts^28^, we identified endothelial cells to acquire the most senescent phenotype (**Suppl. Fig. 5a**). Bulk RNA sequencing data of isolated cardiomyocytes^29^, fibroblasts^28^ and endothelial cells confirmed the up-regulation of senescence marker genes predominantly in aged cardiac endothelial cells (**Suppl. Fig. 5b**). This suggests that endothelial senescence might contribute to neuronal repulsion or death. Indeed, the selective induction of endothelial cell senescence in young animals by endothelial-specific overexpression of progerin^30^ significantly reduced the density of Tuj1 positive nerves compared to wildtype littermates (**Fig. 3h**). Taken together, different models of premature senescence indicate that the induction of (endothelial) senescence is sufficient to induce cardiac sympathetic denervation.

**Fig. 3.**
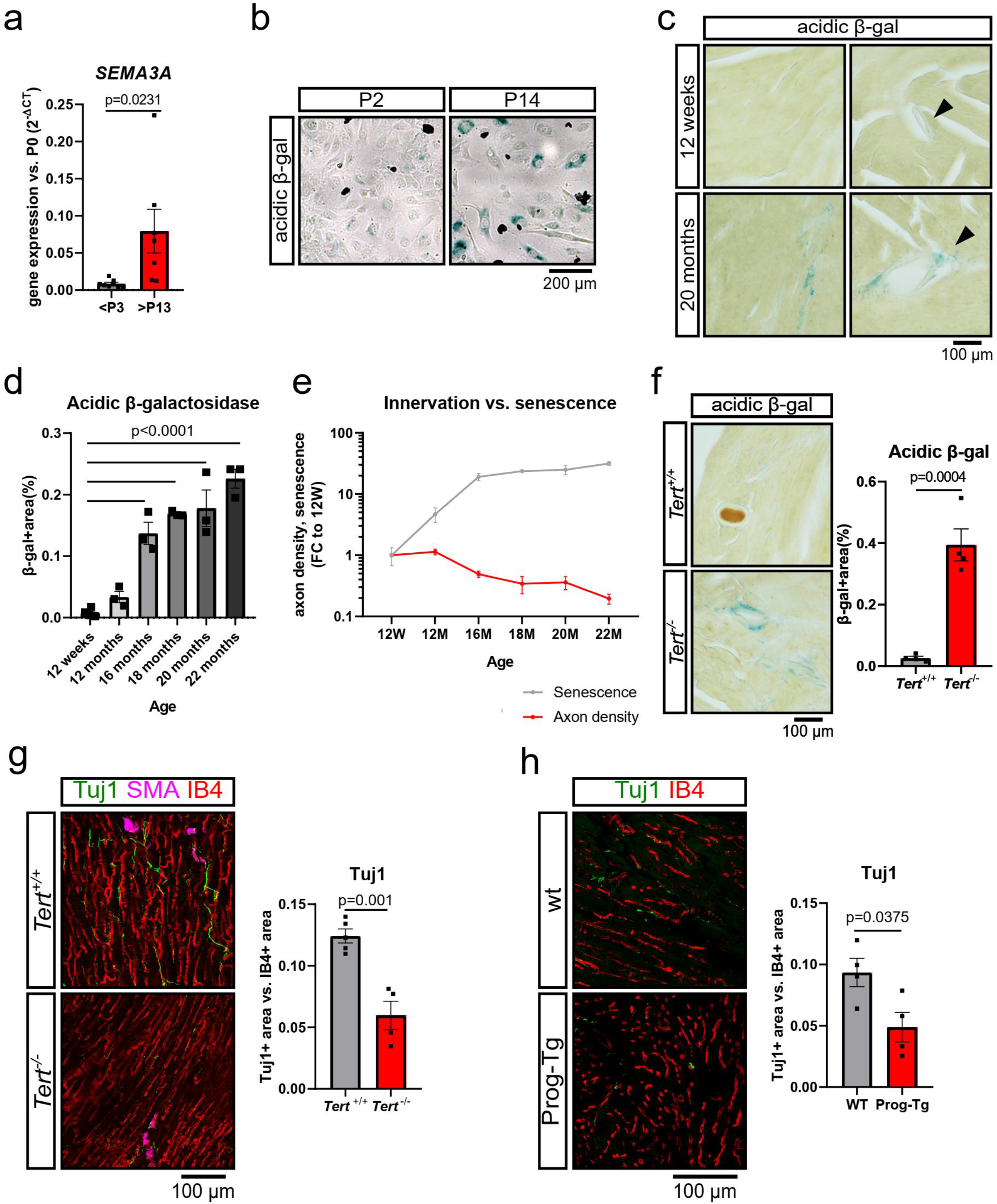
Senescence impairs cardiac innervation. **a** *Sema3a* mRNA expression in short-passage (P2-P3) and long-passage (>P13) HUVEC (n=8 vs. n=7). **b** Representative image of short-passage (P2) and long-passage (P14) HUVEC upon acidic β-galactosidase staining. **c** Representative acidic β-galactosidase stainings on heart sections of male mice at 12 weeks and 20 months old mice (12 weeks: n=6 vs. 20 months: n=3). **d** Quantification of acidic β-galactosidase staining of 12 weeks, 12 months, 16 months, 18 months, 20 months and 22 months of age (12 weeks: n=6 vs. 12 months: n=3, 16 months: n=3, 18 months: n=3, 20 months: n=3, 22 months: n=3). **e** Axon density (Fig. 1e) and acidic β-galactosidase-positive area (Fig. 3c) normed to 12 weeks respectively. **f** Acidic β-galactosidase staining on heart sections of male *Tert*^*−/−*^(4^th^ generation, 10-15 weeks, n=4). **g** Tuj1 staining (green) on heart sections of male *Tert*^*−/−*^(4^th^ generation, 10-15 weeks). IB4 (red) and SMA (magenta) served as counter stain (n=4). **h** Tuj1 staining (green) on heart sections of female and male Prog-Tg mice (28-29 weeks old). IB4 (red) served as counter stain (n=4). Data are shown as mean and error bars indicate the standard error of the mean (SEM). After passing normality test, statistical power was assessed using the unpaired, two-sided t-test.

To determine if interfering with cellular senescence might prevent cardiac denervation in the aged heart, we treated old mice with 5 mg/kg dasatinib and 50 mg/kg quercetin, a combination of senolytics, which was shown to reduce the number of senescent cells by targeting anti-apoptotic pathways and expands life span *in vivo*^31,32^. Treatment was applied via oral gavage to aged mice (18 months) on three consecutive days, every second week for a total duration of two months (**Fig. 4a**). At 2 months after start of the treatment, the number of senescent acidic β-galactosidase-positive cells was significantly lower as compared to placebo-treated controls (**Fig. 4b, c**). Importantly, the reduction in senescent cells was paralleled by a rescue of Tuj1-positive nerves by senolytic treatment (**Fig. 4d**). Consistently, senolytic treatment augmented heart rate variability as assessed by the LF/HF ratio already 2 weeks after start of the treatment (**Fig. 4e, f**). Two months of senolytics treatment further improved the LF/HF ratio in aged mice, and restored the characteristic day-night-rhythm, while old control mice further deteriorated in the autonomous function (**Fig. 4f**; **Suppl. Fig. 6a, b**). These data indicate that senolytics induce a re-innervation of the aging heart, which restores the sympatho-vagal balance. In addition, senolytics treatment improved cardiac function as evidenced by a normalized diastolic function at 4 and 8 weeks of treatment (**Suppl. Fig. 7a, b**) and reduced vulnerability to arrhythmia as assessed by Langendorff-perfused hearts 8 weeks after senolytics treatment (**Suppl. Fig. 8**).

**Fig. 4.**
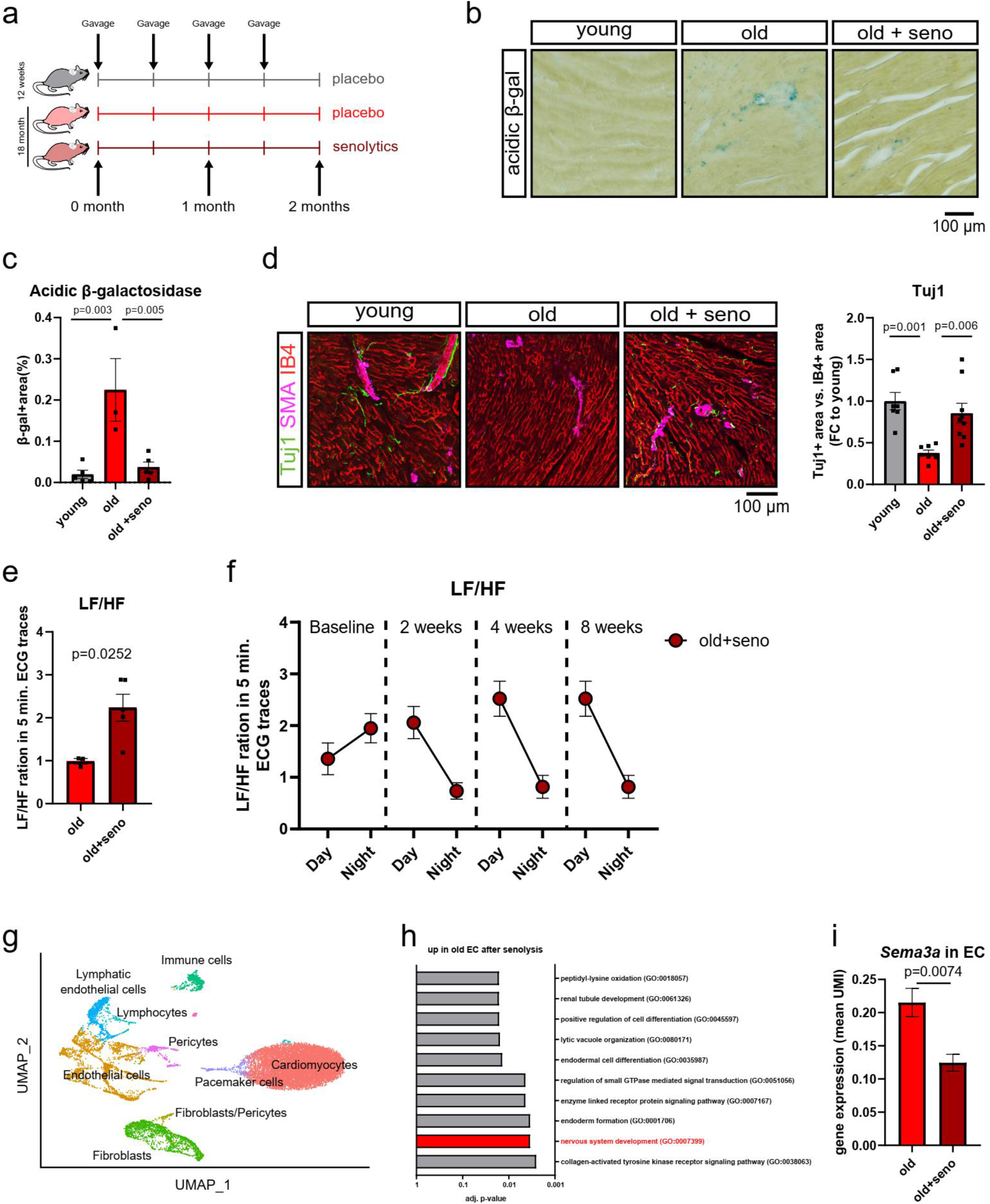
Senolysis rescues axon density in aged mice. **a** Schematic experimental set-up. 18 to 19 months old male mice received a combination of the two senolytics drugs dasatinib and quercetin on three consecutive days, every second week, for a total duration of two months. Young (16 weeks) and old (18 to 19 months) mice receiving the vehicle (placebo) served as control cohorts. **b /c** Acidic β-galactosidase staining on heart sections of male old mice after senolytics treatment vs. heart sections of the respective control groups as described in panel a. Panel c shows the quantification of acidic β-gal positive area (n=5 vs. n=3 vs. n=5). **d** Tuj1 (green), SMA (magenta) and IB4 (red) staining on heart sections of male old mice after senolytics treatment vs. heart sections of the respective control groups as described in panel a (n=7 vs. n=7 vs. n=9). **e** Frequency domain measurement (LF/HF ratio) of 5 min. ECG traces at day time in old mice after two weeks of placebo vs. senolytics treatment (n=3 vs. n=5). **f** Frequency domain measurement (LF/HF ratio) of 5 min. ECG traces at day and night time in old mice before and after 2, 4 and 8 weeks of senolytics treatment (n>5). **g** UMAP visualizing single nuclei RNA sequencing of old male mice after placebo and senolytics treatment (n=3). **h** GO terms analysis of significantly up-regulated genes in endothelial cells upon senolytics treatment. Listed are the top-10 regulated GO terms. **i** *Sema3a* mRNA (mean UMI expression) expression in the endothelial, pericyte and fibroblast clusters of the single nuclei RNA sequencing data shown in panel f. Data are shown as mean and error bars indicate the standard error of the mean (SEM). After passing normality test, statistical power was assessed using the unpaired, two-tailed t-test (d). For comparing multiple groups an ordinary one-way ANOVA with a post-hoc Tukey test was used (b, c). To assess the statistical power of single nuclei RNA sequencing data, a cluster t-test was used (h).

To provide mechanistic insights into how senolytics rescue cardiac innervation, we performed single nuclei RNA sequencing of old mouse hearts treated with senolytics or placebo (**Fig. 4g**). Interestingly, senolytic treatment affected genes associated with “nervous system development” within the top-regulated genes of cardiac endothelial cells (**Fig. 4h**). Importantly, *Sema3a*, which we showed to be de-repressed in the aging heart, was significantly reduced in old heart endothelial cells after senolytic treatment (**Fig. 4i**).

Here, we demonstrate that aging reduces axon density in the heart. Aging induced decline in axon density was associated with reduced miR-145 levels and de-repression of its target, the neuronal repulsive signal Semaphorin-3A, which is well known to induce electrical instability in the heart. Interestingly, induction of cellular senescence, which is a hallmark of aging, was inversely correlated with the onset of axon decline. Targeting senescent cells pharmacologically was sufficient to prevent the decline in axon density and reduced *Sema3a* expression in the aging heart suggesting a key role of senescent cells in cardiac denervation. Senescent cells release numerous secreted factors, termed senescence-associated secretory phenotype (SASP), which profoundly alters the microenvironment in the aging heart. Although neuronal guidance factors have not been reported as general SASPs, Semaphorin-3A is induced in senescent endothelial cells and may represent a specific vascular SASP in the aging heart.

The question which cell type(s) further contributes to the observed effects will need further studies. We demonstrate that the selective induction of pre-mature aging in endothelial cells is sufficient to reduce nerve density. However, we cannot exclude the involvement of other cells in this process. Interestingly, there was a tendency that *Sema3a* was also de-repressed by senolytic treatment in other vascular cells, namely pericytes, whereas e.g. fibroblasts showed very low levels and no *Sema3a* regulation (**Suppl. Fig. 9**). Moreover, neuronal and axonal related pathways were found within the top-25-regulated GO terms in lymphatic endothelial cell and in some fibroblast clusters of mice treated with senolytics (**Suppl. Tab. 1**). These findings indicate that endothelial cells may play a critical role in age-related denervation, but other cells such as pericytes or fibroblasts may contribute as well to the observed phenotype.

Our study additionally demonstrates that senolytic treatment restores vulnerability to arrhythmia, heart rate variability and the circadian rhythm. A decline in heart rate variability is typically observed in the elderly, is indicative of impaired sympathetic and parasympathetic innervation and is associated with increased electrical instability leading to increased overall mortality^11^. Our finding that senolytics normalizes heart rate variability during aging, thus, supports a functional benefit of the treatment.

Innervation is not only important for the control of heart rhythm but nerves were shown to provide important paracrine factors, which for example contribute to cardiac regeneration^12,13^. A decline in nerve density may consequently lead to depletion of such nerve-derived factors influencing the reparative function of the heart as it was demonstrated for myocardial infarction in adult mice^33^. Moreover, in other tissues, nerves interact with immune cells^34^ and can control vascular inflammation^14^. Since inflammation is a hallmark of aging (“inflamaging”), the relation of neuro-immune interactions in the heart may deserve further studies. Together, the presented findings may lay the ground to decipher neuronal cross-talks in the heart and their role in aging.

## Methods

### Laboratory animals

Isogenic male C57Bl/6J wildtype mice were purchased from Janvier (Le Genest SaintIsle, France) and from Charles River (Sulzfeld, Germany). Homozygosity of these inbred mice was controlled by Janvier and Charles River using exome sequencing.

*miR143/145* gene cluster knockout mice were generated as previously described^35^. Male and female miR143/145 gene cluster knockout mice with an C57Bl/6J background and an age between 10 to 15 weeks were used.

Pre-mature senescence was studied in male *Tert*-knockout mice (4^th^ generation, 10 to 15 weeks old) with an C57Bl/6J background as previously described^36,37^.

Endothelial-specific progeria mice were generated as previously described^30^. Male and female mice were use at the age of 28 to 29 weeks.

To obtain hearts, mice were sacrificed via cervical dislocation during isoflurane anesthesia and perfused with cold Hank’s buffered saline solution (HBSS; 14175-053, Invitrogen).

Mice were housed in individually ventilated cages in a specific pathogen-free facility according to national and institutional guidelines for animal care.

### Senolytic treatment

To eliminate senescent cells from aged mice, a combination of the two senolytics drugs dasatinib and quercetin was used as proposed by Xu et al.^38^. In brief, 5 mg/kg dasatinib (SML2589-50MG; Merck) and 50 mg/kg quercetin (Q4951-10G, Sigma-Aldrich) were applied via oral gavage to aged mice (18 months) on three consecutive days, every second week for a total duration of two months. Young (12-16 weeks) and aged (18-19 months) mice receiving the solvent (Phosal 50PG (368315, Lipoid) containing 3.3% ethanol (32221, Sigma-Aldrich) and 10% polyethylene glycol 400 (807485, Merck)) served as control cohorts. Cardiac function was monitored during the experiment using echocardiography and ECG traces as described below. The animal experiment has been conducted as approved by the state of Hessen (animal application number FU/1269).

### Echocardiography

To assess heart function via echocardiography, mice were anaesthetized (2-2.5% isoflurane) and monitored using the Vevo 3100 echocardiography system with the Vevo LAB software (Fujifilm VisualSonics).

### Telemetric ECG measurement

To record long-term ECG traces remotely in awake mice, ETA-F10 transmitters (270-0160-002, DSI) were implanted subcutaneously as described by the provider’s instruction. In brief, buprenorphine (0.1 mg/kg) was injected i.p. to mice 30 minutes before starting the surgery. Then mice were anaesthetized (1.5% isoflurane) and an incision was made on the left anatomical side of the mouse. The transmitter was covered with polymyxin and placed subcutaneously. The electrodes were stitched to the pectoral muscles in Einthoven II position. The wound was closed and mice received metamizol on three consecutive after surgery as post-surgical treatment. The animal experiment has been conducted as approved by the state of Hessen (animal application number FU/1269). ECG traces were recorded and time and frequency domain were analyzed using the software Ponemah 6.

### Assessment of ventricular arrhythmia inducibility

After anaesthesia by isoflurane and cervical dislocation hearts were rapidly excised by opening the thorax and immediately placed in ice-cold buffer solution (modified Krebs-Henseleit solution; mM: NaCl 119, NaHCO_3_ 25, KCL 4.6, KH_2_PO_4_ 1.2, MgSO_4_ 1.1, CaCl_2_ 2.5, C_6_H_12_O_6_8.3 and Na-Pyruvate 2; pH 7.4)^5^. The ascending aorta was pulled over a cannula, hearts were transferred into a Langendorff apparatus and the cannula was rapidly attached to keep no flow time as short as possible. Hearts were electro-mechanical uncoupled by blebbistatin added to the perfusion buffer (5 – 10 µM, Hoelzel Biotech). Perfusion pressure and heart rate were continuously monitored (Powerlab 8/30 & Labchart, ADInstruments). Perfusion flow was manually regulated based on the perfusion pressure (80 – 100 mmHg) using a peristaltic pump (Regalo Masterflex Masterflex, Ismatec)^39^. An octopolar electrophysiology catheter (2.0 F, 0.5 mm electrode spacing; CIBer Mouse, NuMed) was placed in the right ventricle to stimulate the heart and to continuously obtain atrial and ventricular electrograms^5,40^. For equilibration the heart was paced at 600 bpm for 30 minutes. Perfusion buffer was continuously oxygenated using carbogen (95% O_2_/5% CO_2_). Hearts which presented relevant arrhythmias or visible ischemia after equilibration were excluded. To assess susceptibility to ventricular arrhythmias (VA) we used a stimulation protocol based on three maneuvers: (1) Programmed extrastimulation: train of eight S1-stimuli (cycle length (CL) 100 ms) followed by two or three extrastimuli with a decremental S2S3- or S3S4-interval with a stepwise (2 ms) reduction (60 – 20 ms). (2) Miniburststimulation: train of 20 S1-stimuli (CL 100 ms) followed by ten S2-stimuli with a decremental S2-interval with a stepwise (2 ms) reduction (60 – 20 ms). (3) Burststimulation: train of 20 – 100 S1-stimuli with a decremental S1-interval (50 – 10 ms). VAs were classified using an established scoring system^40^.

### Single-nucleus RNA sequencing

To assess the cardiac transcriptome on single nuclear level, nuclei isolation from mouse hearts, single-nuclei separation, library preparation and sequencing were performed as previously described^41^.

### Single-nucleus RNA sequencing data analyses

To analyze single-nucleus RNA sequencing mouse data of senolytic and the control treated mice, the samples were mapped to the mice reference genome (GRCm38) via STARsolo (version 2.7.9) with the parameter “-- soloFeatures GeneFull”. Data ingtegration, normalisation, scaling and UMAP clustering were performed with Seurat (version 4.1.1), according to the developer’s tutorial (https://satijalab.org/seurat/articles/pbmc3k_tutorial.html). After filtering of nuclei based on mitochondrial content (<5%) and genes per nucleus (<2500) a total 13541 single nuclei were analyzed from 6 different samples.

Differential expressed genes were tested using the FindAllMarkers function with the statistical test bimod in the Seurat package. Genes with adjusted p-values <0.05 were considered as differential expressed genes.

### Whole mount immunofluorescence staining

After sacrificing mice, hearts were perfused with 4% PFA (28908, ThermoFisher Scientific) in PBS, harvested and incubated in 4% PFA for 4h at 4°C. Hearts were washed trice with PBS for 5 minutes and bleached overnight at room temperature using DMSO (A994.2, Carl Roth GmbH & Co. KG) and H_2_O_2_ (8070.2, Carl Roth GmbH & Co. KG) diluted 1:1:4 (vol/vol/vol) in PBS. Hearts were washed trice with PBS for 20 minutes each followed by antigen retrieval by incubating whole hearts in retrieval buffer (4% SDS (CN30.3, Carl Roth GmbH & Co. KG) and 200 mM boric acid (191411, MP Biomedicals)) for 1h at room temperature followed by overnight incubation at 54°C. Hearts were again washed trice in PBT (0.2% Triton X-100 in PBS) for 1h each and incubated in blocking solution (10% FBS (4133, Invitrogen), 1% BSA (A7030-10G, Merck), 5% donkey serum (017-000-121, Jackson Immuno) in PBT) for 1h at room temperature. Rabbit anti-Tuj1 antibody (ab18207, Abcam) was diluted 1:100 in blocking solution and incubated with the hearts for 3 days at room temperature. Hearts were then washed trice in PBS for 20 minutes and incubated with the secondary antibody (donkey anti-rabbit antibody conjugated to Alexa 555; A-31572, Invitrogen) that was diluted 1:100 for at least 2 days at room temperature. Hearts were again washed trice for 20 minutes in PBS and embedded in agarose (9012-36-6, Carl Roth GmbH & Co. KG). Hearts were dehydrated at room temperature in an ascending methanol series (30%, 50%, 75%, 30 minutes each) and incubated twice in 100% methanol at room temperature for 30 minutes each. Hearts were washed twice in ECI (112372, Sigma-Aldrich) for 5 minutes and cleared by incubating in 80% ECI and 20% PEGM (447943, Sigma-Aldrich) for 30 minutes at room temperature.

Whole hearts were assessed histologically using a light sheet microscope (Ultramicroscope II, LaVision BioTec, Bielefeld, Germany). Excitation was performed at 470/40 nm and emission 525/50 nm (autofluorescence tissue), excitation 545/30 nm and emission 595/40 nm(Tuj). Main laser power 95% and software laser power for 470/40 95% and 525/50 35%. Step size was set to 5 µm. Exposure time was 300 ms, 6,3× magnification (10× zoom body + 0.63× Objective). Sheet width 60%; Sheet NA 4,05um; two sided scan. Pictures were taken with a Neo 5.5 (3-tap) sCOMs Camera (Andor, Mod. No.: DC-152q-C00-FI). Images were analyzed using the Imaris software, version 9.

### Immunofluorescence staining of cryopreserved heart sections

After sacrificing mice, hearts were flushed with cold HBSS and fixed in PBS containing 4% PFA (28908, ThermoFisher Scientific). After overnight incubation at 4°C, hearts were washed three times for 10 minutes in PBS. To cryopreserve cardiac tissues, three consecutive overnight washes in PBS containing increasing concentrations of sucrose (10%, 20%, 30%; S0389, Sigma-Aldrich) were applied at 4°C. Tissues were embedded in PBS containing 15% sucrose, 8% gelatin (G1890, Sigma-Aldrich), and 1% polyvinylpyrrolidone (P5288, Sigma-Aldrich). After the embedding solution was solidified, tissues were stored at −80°C. Hearts were sectioned at 50 µm-thickness using a cryostate (Leica CM3050 S). Sections were placed on adhesive glass slides (10149870, ThermoFisher Scientific) and stored at −20°C until use.

For immunofluorescence staining, cryo sections were brought to room temperature and re-hydrated in PBS (twice for 5 minutes). To permeabilize the tissue, sections incubated with PBS containing 0.3% Triton X-100 three times for 10 minutes and were blocked in PBS containing 0.1% Triton X-100, 3% BSA (A7030-10G, Merck) and 5% donkey serum (ab7475, Abcam) for 1h at room temperature. Primary antibodies were diluted in blocking solution and incubated with the sections overnight at 4°C. Sections were then washed three times for 5 minutes in PBS and incubated for 1h at room temperature with the respective secondary donkey antibodies that were diluted in PBS containing 0.1% Triton X-100. Nuclei were stained with DAPI (6335.1, Carl Roth GmbH & Co. KG) that was diluted 1:1000 in 0.1% Triton X-100. After washing trice in PBS for 5 minutes, slides were mounted with Fluoromount-G™ (00-4958-02, Invitrogen). Sections were histologically assessed using the Leica Stellaris confocal microscope and the LASX software.

### Immunofluorescence staining of paraffin heart sections

To assess hearts histologically on paraffin sections, hearts were processed and embedded as previously described^28^. To immunolabel paraffin section, slides incubated for 1h at 60°C and were deparaffinized twice with xylene for 10 minutes and an ethanol series of 100%, 95%, 80%, 70%, and 50% ethanol (5 minutes each step). Sections were washed in water for 5 minutes and were boiled in 0.01 M citrate buffer (pH = 6) for 90 seconds. Slides were then washed for 5 minutes with PBS and blocked in PBS containing 0.1% Triton X-100, 3% BSA (A7030-10G, Merck) and 5% donkey serum (ab7475, Abcam) for 1h at room temperature. Primary antibodies were diluted in blocking solution and incubated with the sections overnight at 4°C. Sections were then washed three times for 5 minutes in PBS and incubated for 1h at room temperature with the respective secondary donkey antibodies that were diluted in PBS containing 0.1% Triton X-100. Nuclei were stained with DAPI (6335.1, Carl Roth GmbH & Co. KG) that was diluted 1:1000 in 0.1% Triton X-100. After washing trice in PBS for 5 minutes, slides were mounted with Fluoromount-G™ (00-4958-02, Invitrogen). Sections were histologically assessed using the Leica Stellaris confocal microscope and the LASX software.

### Antibodies

Following primary antibodies have been uses:

Rb anti-Tuj1 (1:100, ab18207, Abcam), Rb anti-tyroxine hydroxylase (1:100, AB152, Merck), Gt anti-Choline Acetyltransferase (1:100, AB144P, Merck), Gt anti-calcitonin gene-related peptide (1:100, ab36001, Abcam), Ms anti-α-Smooth Muscle - Cy3™ (1:200, C6198-2ML, Sigma-Aldrich), Rb anti-Semaphorin-3A (1:100, ab23393, Abcam) and GSL I - isolectin B4 (biotinylated; 1:25, VEC-B-1205, Biozol).

Following secondary antibodies have been used:

Donkey anti-mouse IgG Alexa Fluor 647 (1:200, A-31571, Invitrogen), Donkey anti-rabbit IgG Alexa Fluor 555 (1:200, A-31572, Invitrogen), Donkey anti-rabbit IgG Alexa Fluor 488 (1:200, A-21206, Invitrogen), Donkey anti-Goat IgG Alexa Fluor 555 (1:200, A-21432, Invitrogen), Donkey anti-Goat IgG Alexa Fluor 647 (1:200, A-21447, Invitrogen), Streptavidin, Alexa Fluor™ 405 (1:200, S32351, Invitrogen) and Streptavidin, Alexa Fluor™ 647 (1:200, S32357, Invitrogen).

### Quantification of immunofluorescence images

To analyze and quantify immunofluorescence images, the stained area was determined and normalized to IB4- or DAPI-positive area using the software Volocity 7 by Quorum Technologies Inc.

### Acidic beta-galactosidase staining

Acidic beta-galactosidase positive cells were visualized on cryopreserved heart sections and *in vitro* using the Senescence β-Galactosidase Staining kit (9860, CST) according to the manufacturer’s instruction. β-galactosidase-positive areas were quantified using ImageJ.

### Endothelial cell isolation from murine hearts

Cardiac endothelial cells were isolated from young (12 weeks) and old (20 months) mice. Under isoflurane anesthesia, mice were sacrificed and hearts were flushed with HBSS. The hearts were harvested, dissected in small pieces, transferred into a C-tube (130-096-334, Miltenyi Biotec) and incubated in HBSS, containing 600 U/mL collagenase type II (354236, Corning), at 37°C and 5% CO_2_ in a humidified atmosphere. After 30, 20 and 10 minutes of incubation, tissue particles were further dissected using the GentleMACS Dissociator (Miltenyi BioTec) with the pre-set program m_neoheart_01_01. Collagenase digestion was stopped with 500 µL fetal bovine serum (4133, Invitrogen). Cell suspension was applied on a 200 µm cell strainer (43-50200-03, pluri-Select), centrifuged at 80x g and 4°C for 1 minute to deplete cardiomocytes and applied on a 70 µm cell strainer (43-50070-03, pluri-Select). Cells were washed twice with HBSS containing 0.5% bovine serum albumin (T844.3, Carl Roth GmbH & Co. KG) and 2 mM EDTA (A4892, AppliChem) (referred to as wash buffer in the following) by centrifugation (300x g at 4°C for 10 minutes). Endothelial cells were isolated using rat anti-mouse CD144 antibodies (555289, BD Bioscience) and magnetic sheep anti-rat dynabeads (11035, Life Technologies). During tissue dissection, anti-CD144 antibody-bead mixture was prepared by washing 25 µL dynabeads twice with wash buffer and re-suspending them in 400 µL wash buffer. 1 µL of antibodies were added to the beads, incubated for 1h at room temperature and were washed trice with wash buffer. Antibody-bead mixture was re-suspended in 1000 µL wash buffer, added to the cardiac cell pellet and incubated for 40 minutes on a turning wheel. Cells were washed trice on a magnetic rack using 1000 µL of wash buffer and lysed with 700 µL of Qiazol.

### RNA isolation

Total RNA was purified from cultured and isolated cells by using the miRNeasy Mini kits (217004, Qiagen), combined with on-column DNase digestion (DNase Set, 79254, Qiagen) as described in the manufacturer’s instruction. To isolate RNA from solid hearts, tissue was combined with 700 µL Qiazol and ¼” ceramic spheres and were homogenized three times for 20 seconds (4 m/s). RNA isolation was then performed using the miRNeasy Mini kit (217004, Qiagen). The RNA concentration was determined by measuring absorption at 260 nm and 280 nm with the NanoDrop ND 2000-spectrophotometer (PeqLab).

### cDNA synthesis and quantitative PCR

To quantify mRNA expression by qPCR, 100 ng to 1 μg of total RNA was reverse-transcribed using the reverse transcriptase M-MLV (28025013, ThermoFisher Scientific) and assessed using the SYBR™ Green PCR Master Mix (4385617, Applied Biosystems) as previously described^28^. The primers were customized and purchased from Sigma-Aldrich (now Merck): human Rpl0 fw (TCGACAATGGCAGCATCTAC); human Rpl0 rev (ATCCGTCTCCACAGACAAGG); human SEMA3A fw (TGTTGGGACCGTTCTTAAAGTAGT); human SEMA3A rev (TAGTTGTTGCTGCTTAGTGGAAAG).

### Bulk RNA sequencing

Library preparation and whole transcriptome analysis of isolated cardiac endothelial cells were performed as previously described^28^.

### Gene ontology term analysis

Gene ontology (GO-term) analyses were performed using the Enrichr online platform (ontology category GO Biological Process 2021; https://maayanlab.cloud/Enrichr/) by assessing significantly regulated genes of logFC > 1 and logFC < −1.

### Micro Array

To assess micro-RNA expression in whole young and old mouse hearts, a published data set was used^42^.

### Cell Culture

Human umbilical cord vein endothelial cells (HUVEC, CC-2935) were purchased from Lonza and cultured with endothelial basal medium (EBM, CC-3156, Lonza) supplemented with 10 % FBS (4133, Invitrogen), Amphotericin-B (CC-4081C, Lonza), ascorbic acid (CC-4116C, Lonza), bovine brain extract (CC-4092C, Lonza), endothelial growth factor (CC-4017C, Lonza), gentamycin sulfate (CC-4081C, Lonza), and hydrocortisone (CC-4035C, Lonza) at 37°C and 5% CO_2_, with humidified atmosphere. Short passage HUVEC (P2, P3) were used for *in vitro* studies. To resemble cellular senescence, HUVEC were cultured until at least passage 13 and were controlled by acidic beta-galactosidase staining.

### Transfection experiments

70,000 HUVEC were seeded per well of a 12-well plate (665180, Greiner Bio-One GmbH) and rested for one day at 37°C and 5% CO_2_. Cells were transfected using the Lipofectamine RNAiMax (13778150, Invitrogen) according to the manufacturer’s protocol. Predesigned miR-145-5p precursors (4464066 (ID: MC11480), Ambion by Life Technologies) were used in a final concentration of 10 nM. Cells were cultured for 48h after transfection.

### Statistical Analysis

Data are expressed as mean and error bars indicate the standard error of the mean (SEM). Normality distribution was assessed by using the Kolmogorov-Smirnov or Shapiro-Wilk normality test. For comparing two groups of Gaussian distributed data, statistical power was determined using the unpaired, two-sided student’s t-test. To compare more than two groups, an ordinary one-way ANOVA with a post-hoc Tukey comparison was used.

## Data availability

All data are available within the article, within the supplemental material or from the corresponding author on reasonable request. Transcriptomic data are available from the cited publications or at the GEO (accession code will be provided as soon as the manuscript is accepted for publication).

## Acknowledgement

The study is supported by the German Research Foundation (SFB1366, Project B4; TRR267, Project B3 and the Cluster of Excellence Cardiopulmonary Institute Exc2026/1), the German Center for Cardiovascular Research (DZHK Shared Expertise (B22-014 SE)), the Dr. Rolf-M.- Schwiete Foundation (2021-002) and the European Research Council.

## Author contributions

JUGW and SD planned the project and wrote the manuscript. JUGW, LMK, JP conducted the majority of experiments. SG performed bulk RNA sequencing. LST and DJ performed bioinformatics. WTA performed snRNA sequencing. KAS and MC contributed to transcriptomic and histological analyses. AF, PFM and RPB contributed to telemetry studies. KS, GKB, MMR and GL contributed to histology and whole mount staining. SC, DS, SA, EA, NK, MK and CM contributed to electrophysiology and telemetry studies. GL, CM and AMZ provided conceptual input. TBo and TBr provided miR145 mice. CB provided Tert mice. EN and SOM provided Prog-tg mice.

---

**Suppl. Fig. 1.**
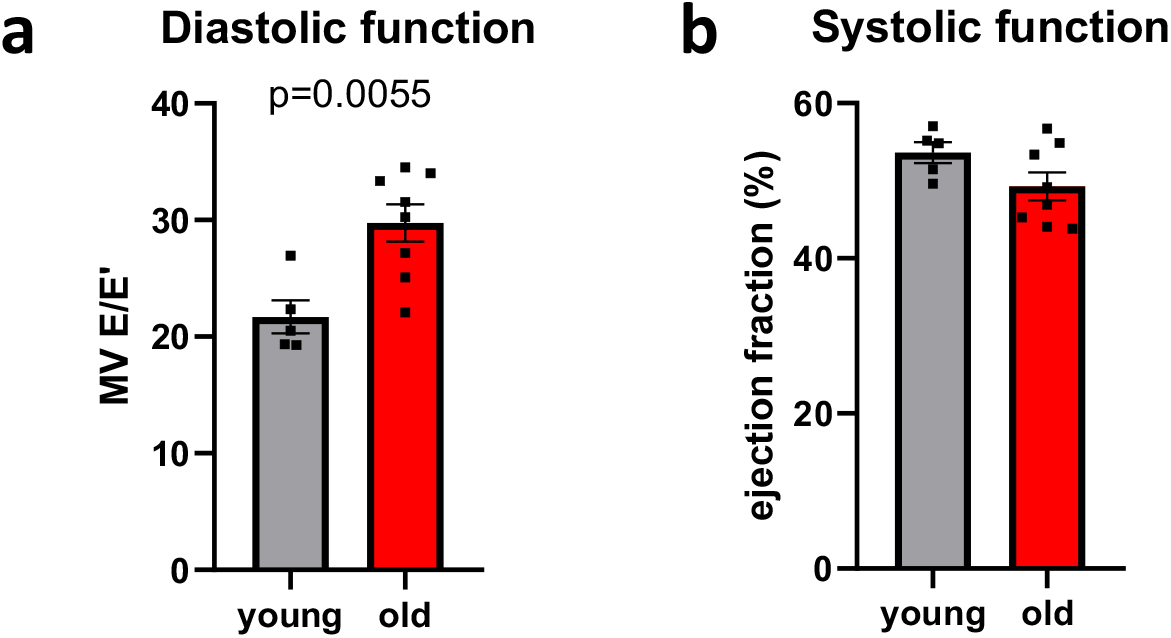
Echocardiography in young vs. old mice. **a** Diastolic function was assessed as MV E/E’ in male young (12 weeks) vs. old (18 months) mice (n=5 vs. n=8). **b** Systolic function was determined by ejection fraction (EF %) in male young (12 weeks) vs. old (18 months) mice (n=5 vs. n=8). Data are shown as mean and error bars indicate the standard error of the mean (SEM). After passing normality test, statistical power was assessed using the unpaired, two-tailed t-test.

**Suppl. Fig. 2.**
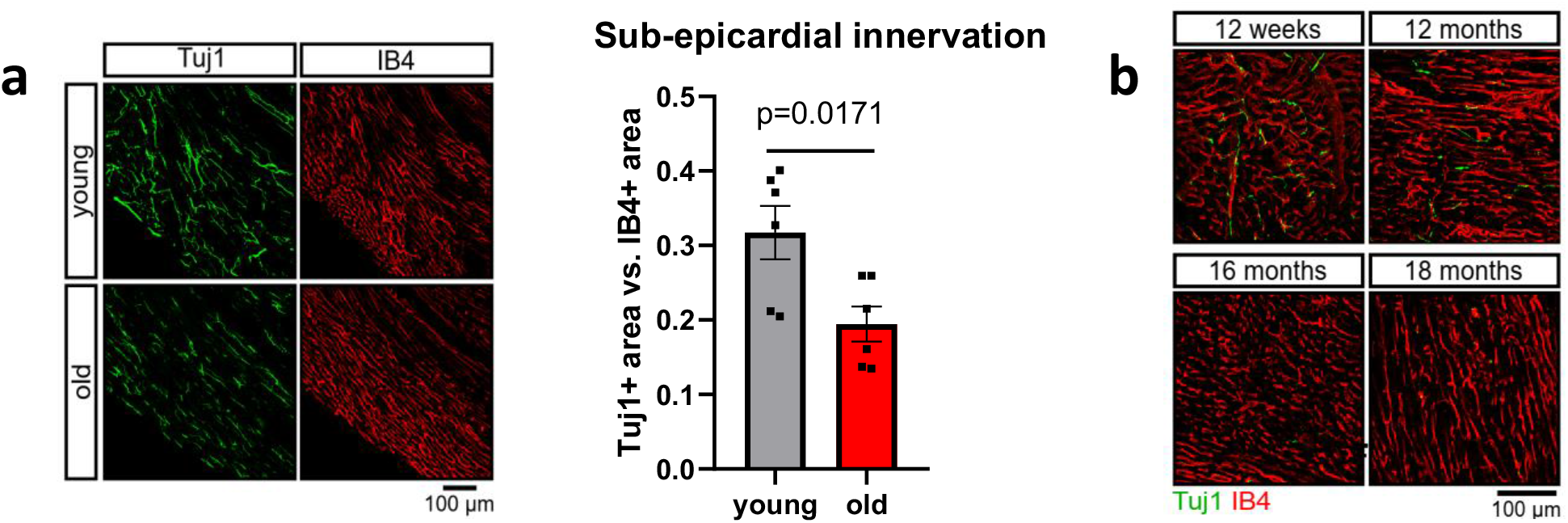
Axon density in aged left ventricles. **a** Histological assessment of the sub-epicardial innervation in left ventricles of young (12 weeks) vs. old (18 months) mice. Axon density was assessed via Tuj1 (green) staining and normalized to IB4 (red) (n=6). **b** Left-ventricular innervation (Tuj1=green) vs. IB4 (red) at 12 weeks, 12 months, 16 months, 18 months, 20 months and 22 months of age (n=12 vs. n=3, n=3, n=8, n=4, n=3).

**Suppl. Fig. 3.**
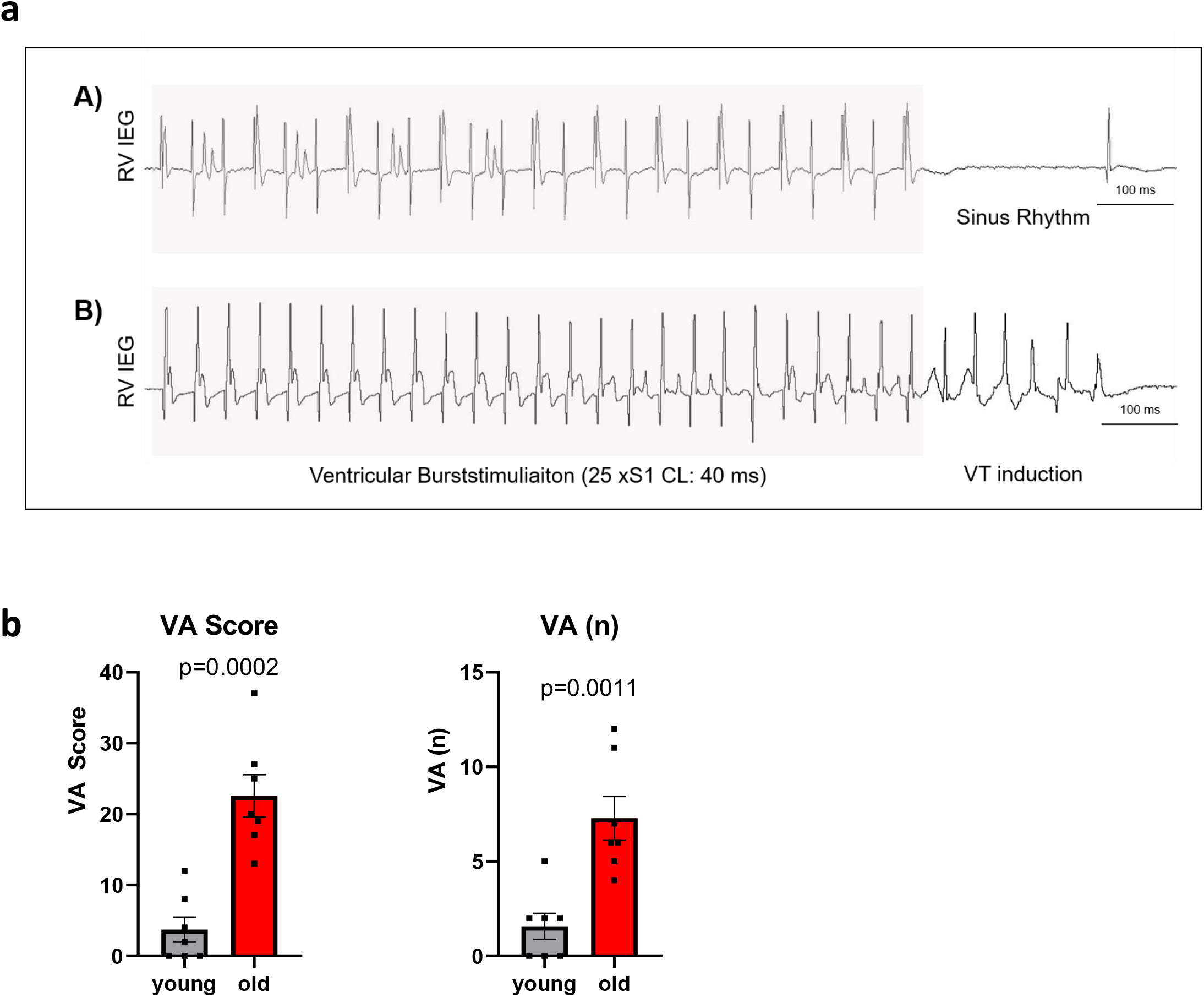
Susceptibility to ventricular arrhythmias is increased in aged hearts. **a** Exemplary right ventricular intracardiac electrograms (RV IEG) of ventricular burststimulation (VBS) maneuvers (grey highlighted) in a young (A) and aged (B) heart ex vivo. In the young heart the VBS does not result in continuous ventricular arrhythmia whereas VBS induces a ventricular tachycardia (VT) in the aged heart. **b** Quantification of ventricular arrhythmias (n=7). Data are shown as mean and standard errors indicate SEM. Statistical power was assessed using unpaired, two-tailed t-test.

**Suppl. Fig. 4.**
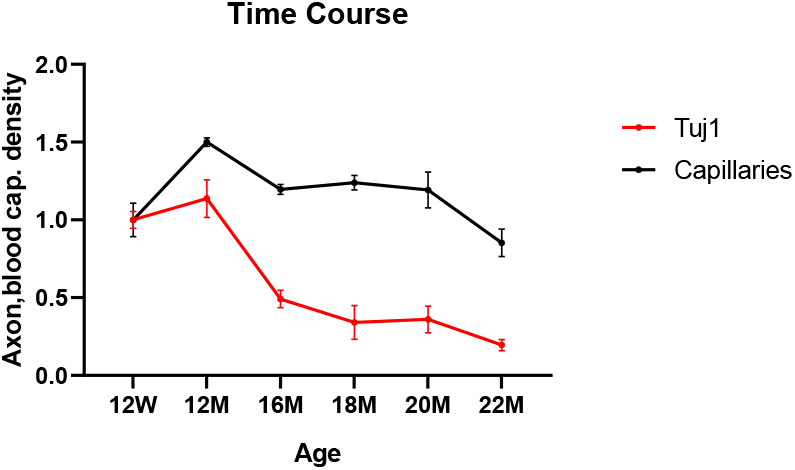
Time course study of axon and capillary density. Axon (Tuj1) and capillary density (IB4) in left ventricles of mice at 12 weeks, 12 months, 16 months, 18 months, 20 months and 22 months of age (n=3).

**Suppl. Fig. 5.**
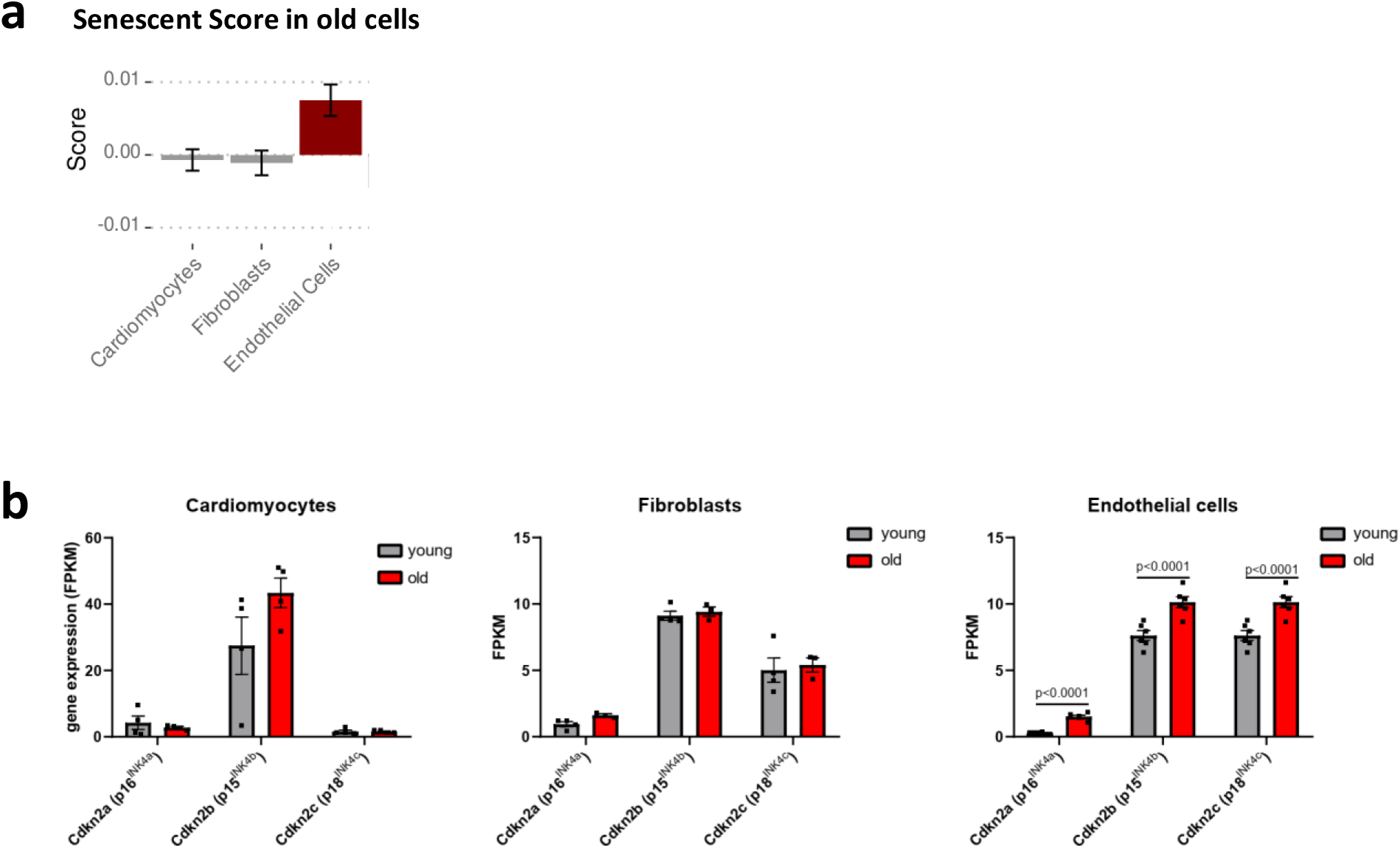
Identifying cellular senescence in old cardiac cells. **a** Senescent score after Kiss et al.,^27^ applied on clusters of cardiomyocytes, fibroblast and endothelial cell derived from published single nuclei RNA sequencing data from young vs. old mouse hearts^28^ (n=3). **b** Senescence marker gene expression in bulk RNA sequencing data of isolated cardiomyocytes^29^ (n=4), fibroblasts^28^(n=4) and endothelial cells (n=6). Data are shown as mean and error bars indicate the standard error of the mean (SEM). After passing normality test, statistical power was assessed using the unpaired, two-tailed t-test.

**Suppl. Fig. 6.**
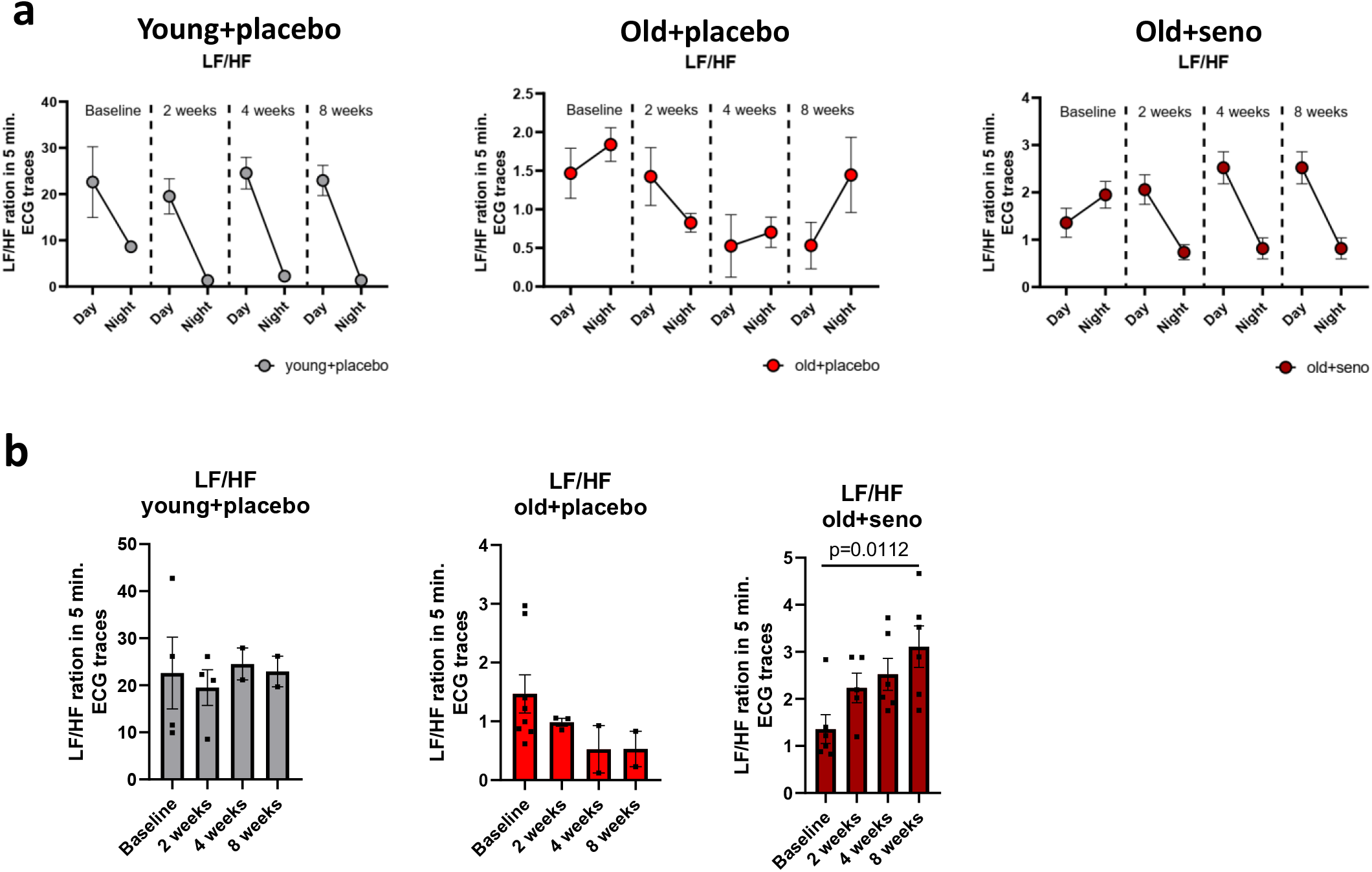
Heart rate variability analyses in old mice upon senolysis. **a/b** Frequency domain measurement (LF/HF ratio) of 5 min. ECG traces at day and night time (a) and statistical analysis of LF/HF ratio at day time only (b). Young placebo (16 weeks), old placebo (19 months) and old senolytics (19 months) mice were analyzed before and after 2 (young-placebo: n=4; old-placebo n=3; old-senolytics: n=5), 4 (young-placebo: n=2; old-placebo n=2; old-senolytics: n=6) and 8 weeks (young-placebo: n=2; old-placebo n=2; old-senolytics: n=6) of treatment. Data are shown as mean and error bars indicate the standard error of the mean (SEM). After passing normality test, statistical power was assessed using an ordinary one-way ANOVA with a post-hoc Tukey test was used (b, old+senolytics).

**Suppl. Fig. 7.**
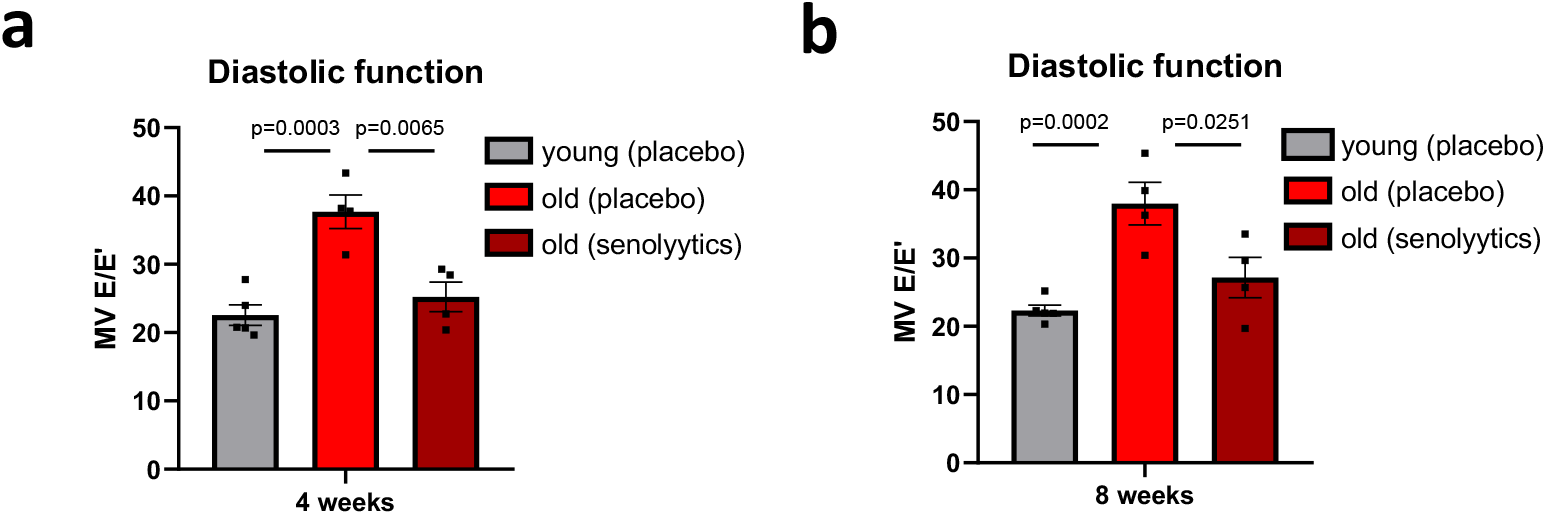
Echocardiography analyses in young and old mice upon senolysis. **a** Diastolic function was assessed as MV E/E’ in male young (12 weeks) vs. old (18 months) mice after 4 weeks of placebo / senolytics treatment (n=5 vs. n=4 vs. n=4). **b** Diastolic function was assessed as MV E/E’ in male young (12 weeks) vs. old (18 months) mice after 8 weeks of placebo or senolytics treatment (n=5 vs. n=4 vs. n=4). Data are shown as mean and error bars indicate the standard error of the mean (SEM). After passing normality test, statistical power was assessed using an ordinary one-way ANOVA with a post-hoc Tukey test was used.

**Suppl. Fig. 8.**
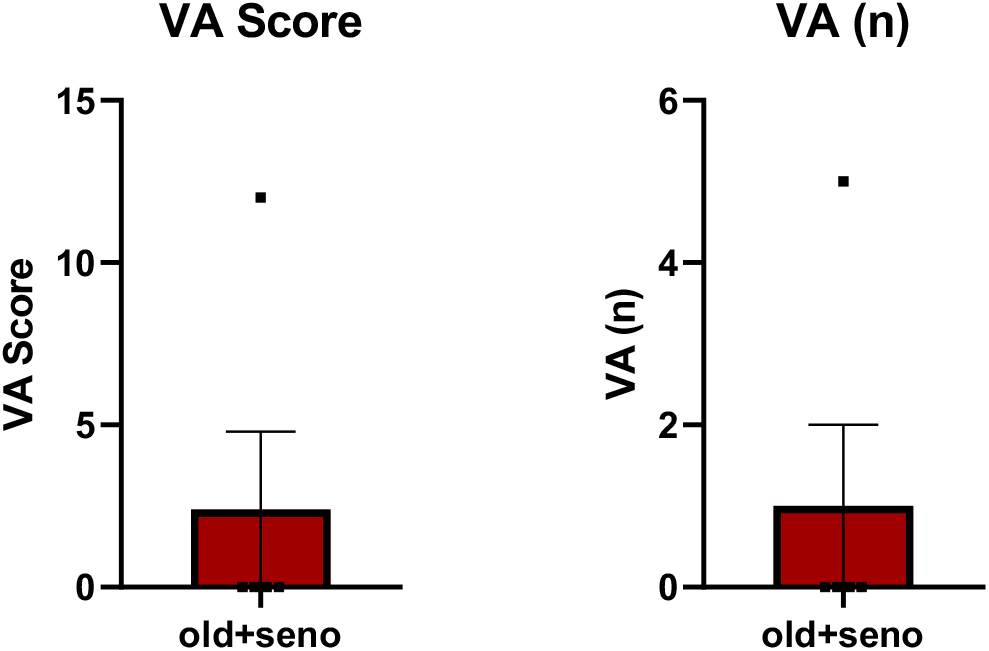
Susceptibility to ventricular arrhythmias is decreased in aged hearts after senolysis. Quantification of ventricular arrhythmias in old hearts, two months after senolytics drug administration (n=5).

**Suppl. Fig. 9.**
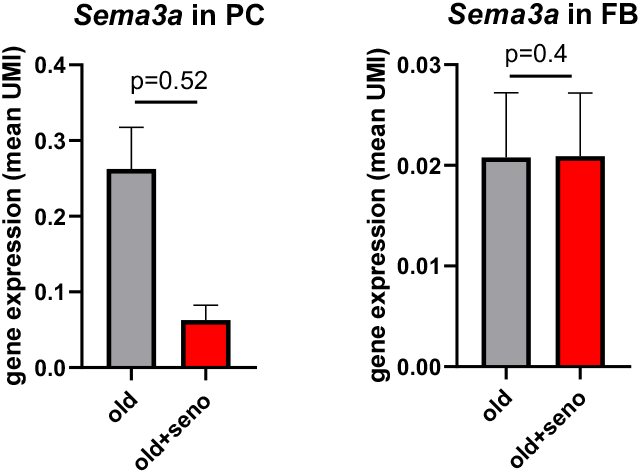
Sema3a expression on single nuclei level. Mean UMI expression showing *Sema3a* expression in pericytes (PC) and fibroblasts (FB) in old vs. old senolytics mouse hearts (n=3). Data are expressed as mean and error bars indicate standard error of the mean. Statistical power was assessed using a clustered t-test.

**Suppl. Tab. 1:** Top 25 GO terms which are up- and down-regulated in endothelial cells (EC), lymphatic endothelial cells (LymphEC), fibroblasts (FB) and pericytes (PC) in aged hearts upon senolytics treatment. Data are received from snRBA seq.

|  | Term | P-value | Adj. P-value |
| --- | --- | --- | --- |
| GO terms, up-regulated in aged cardiac EC upon senolytics | extracellular matrix organization (GO:0030198) | 2,71E-05 | 6,50E-02 |
|  | extracellular structure organization (GO:0043062) | 6,50E-06 | 6,50E-02 |
|  | external encapsulating structure organization (GO:0045229) | 7,59E-05 | 6,50E-02 |
|  | collagen fibril organization (GO:0030199) | 7,09E-01 | 4,55E+02 |
|  | supramolecular fiber organization (GO:0097435) | 2,94E+02 | 1,51E+06 |
|  | collagen-activated signaling pathway (GO:0038065) | 7,20E+08 | 2,79E+11 |
|  | transmembrane receptor protein tyrosine kinase signaling pathway (GO:0007169) | 7,60E+08 | 2,79E+11 |
|  | negative regulation of multicellular organismal process (GO:0051241) | 2,31E+09 | 7,42E+11 |
|  | collagen-activated tyrosine kinase receptor signaling pathway (GO:0038063) | 8,99E+09 | 2,57E-03 |
|  | endoderm formation (GO:0001706) | 1,48E+11 | 3,47E-03 |
|  | nervous system development (GO:0007399) | 1,49E+11 | 3,47E-03 |
|  | enzyme linked receptor protein signaling pathway (GO:0007167) | 2,06E+11 | 4,39E-03 |
|  | regulation of small GTPase mediated signal transduction (GO:0051056) | 2,22E+10 | 4,39E-03 |
|  | endodermal cell differentiation (GO:0035987) | 7,63E+10 | 1,40E-02 |
|  | lytic vacuole organization (GO:0080171) | 9,14E+10 | 1,56E-02 |
|  | positive regulation of cell differentiation (GO:0045597) | 1,01E+12 | 1,63E-02 |
|  | renal tubule development (GO:0061326) | 1,15E+12 | 1,64E-02 |
|  | peptidyl-lysine oxidation (GO:0018057) | 1,15E+12 | 1,64E-02 |
|  | actin filament organization (GO:0007015) | 2,03E+12 | 2,64E-02 |
|  | lysosome organization (GO:0007040) | 2,07E+11 | 2,64E-02 |
|  | aorta development (GO:0035904) | 2,26E+11 | 2,64E-02 |
|  | retinal ganglion cell axon guidance (GO:0031290) | 2,27E+11 | 2,64E-02 |
|  | regulation of actin cytoskeleton organization (GO:0032956) | 2,38E+12 | 2,66E-02 |
|  | positive regulation of phosphorylation (GO:0042327) | 2,57E+12 | 2,75E-02 |
|  | inflammatory response (GO:0006954) | 2,86E+10 | 2,94E-02 |
| GO terms, down-regulated in aged cardiac EC upon senolytics | phosphorylation (GO:0016310) | 3,86E+09 | 3,37E-03 |
|  | negative regulation of cellular macromolecule biosynthetic process (GO:2000113) | 5,71E+10 | 2,49E-02 |
|  | protein phosphorylation (GO:0006468) | 1,46E+12 | 3,16E-02 |
|  | entrainment of circadian clock by photoperiod (GO:0043153) | 1,62E+12 | 3,16E-02 |
|  | photoperiodism (GO:0009648) | 1,81E+12 | 3,16E-02 |
|  | positive regulation of apoptotic process (GO:0043065) | 2,24E+12 | 3,26E-02 |
|  | cellular response to interleukin-21 (GO:0098757) | 4,25E+11 | 4,64E-02 |
|  | interleukin-21-mediated signaling pathway (GO:0038114) | 4,25E+11 | 4,64E-02 |
|  | cellular response to interleukin-9 (GO:0071355) | 5,45E+10 | 4,76E-02 |
|  | interleukin-9-mediated signaling pathway (GO:0038113) | 5,45E+10 | 4,76E-02 |
|  | cellular response to cytokine stimulus (GO:0071345) | 6,26E+11 | 4,97E-02 |
|  | interleukin-35-mediated signaling pathway (GO:0070757) | 8,28E+11 | 5,56E-02 |
|  | positive regulation of posttranscriptional gene silencing (GO:0060148) | 8,28E+11 | 5,56E-02 |
|  | cellular response to mechanical stimulus (GO:0071260) | 9,59E+11 | 5,98E-02 |
|  | positive regulation of fat cell differentiation (GO:0045600) | 1,08E-03 | 6,00E-02 |
|  | cellular response to interleukin-15 (GO:0071350) | 1,17E-03 | 6,00E-02 |
|  | interleukin-15-mediated signaling pathway (GO:0035723) | 1,17E-03 | 6,00E-02 |
|  | growth hormone receptor signaling pathway via JAK-STAT (GO:0060397) | 1,36E-03 | 6,59E-02 |
|  | cellular response to estrogen stimulus (GO:0071391) | 1,56E-03 | 6,83E-02 |
|  | interleukin-27-mediated signaling pathway (GO:0070106) | 1,56E-03 | 6,83E-02 |
|  | negative regulation of gene expression (GO:0010629) | 1,72E-03 | 7,01E-02 |
|  | regulation of rRNA processing (GO:2000232) | 1,78E-03 | 7,01E-02 |
|  | positive regulation of cytokine production involved in inflammatory response (GO:1900017) | 2,02E-03 | 7,01E-02 |
|  | regulation of protein-containing complex disassembly (GO:0043244) | 2,26E-03 | 7,01E-02 |
|  | positive regulation of cell death (GO:0010942) | 2,27E-03 | 7,01E-02 |
| GO terms, up-regulated in aged cardiac lymphEC upon senolytics | glycolipid metabolic process (GO:0006664) | 5,67E+10 | 1,18E-01 |
|  | regulation of macrophage migration (GO:1905521) | 8,56E+10 | 1,18E-01 |
|  | actin filament-based transport (GO:0099515) | 1,73E+12 | 1,42E-01 |
|  | regulation of cell migration (GO:0030334) | 2,05E+12 | 1,42E-01 |
|  | positive regulation of cell migration (GO:0030335) | 2,73E+11 | 1,42E-01 |
|  | negative regulation of actin filament depolymerization (GO:0030835) | 3,35E+11 | 1,42E-01 |
|  | positive regulation of signal transduction (GO:0009967) | 3,65E+12 | 1,42E-01 |
|  | positive regulation of endothelial cell migration (GO:0010595) | 4,11E+12 | 1,42E-01 |
|  | endothelial cell migration (GO:0043542) | 4,87E+11 | 1,49E-01 |
|  | positive regulation of cell motility (GO:2000147) | 7,46E+11 | 2,06E-01 |
|  | regulation of Cdc42 protein signal transduction (GO:0032489) | 9,03E+11 | 2,25E-01 |
|  | endothelial cell proliferation (GO:0001935) | 1,06E-03 | 2,25E-01 |
|  | T cell migration (GO:0072678) | 1,06E-03 | 2,25E-01 |
|  | regulation of small GTPase mediated signal transduction (GO:0051056) | 1,22E-03 | 2,40E-01 |
|  | vesicle transport along actin filament (GO:0030050) | 1,31E-03 | 2,42E-01 |
|  | positive regulation of epithelial cell proliferation (GO:0050679) | 1,47E-03 | 2,54E-01 |
|  | ganglioside metabolic process (GO:0001573) | 1,60E-03 | 2,61E-01 |
|  | cellular response to vascular endothelial growth factor stimulus (GO:0035924) | 1,78E-03 | 2,67E-01 |
|  | regulation of lipid biosynthetic process (GO:0046890) | 2,04E-03 | 2,67E-01 |
|  | semaphorin-plexin signaling pathway (GO:0071526) | 2,04E-03 | 2,67E-01 |
|  | negative regulation of BMP signaling pathway (GO:0030514) | 2,28E-03 | 2,67E-01 |
|  | regulation of endothelial cell migration (GO:0010594) | 2,29E-03 | 2,67E-01 |
|  | negative regulation of voltage-gated potassium channel activity (GO:1903817) | 2,51E-03 | 2,67E-01 |
|  | phagosome maturation (GO:0090382) | 2,62E-03 | 2,67E-01 |
|  | regulation of actin cytoskeleton organization (GO:0032956) | 2,64E-03 | 2,67E-01 |
| GO terms, down-regulated in aged cardiac lymphEC upon senolytics | ncRNA processing (GO:0034470) | 1,55E+12 | 1,00E-01 |
|  | positive regulation of RNA polymerase II transcription preinitiation complex assembly (GO:0045899) | 3,28E+12 | 1,00E-01 |
|  | regulation of G2/M transition of mitotic cell cycle (GO:0010389) | 3,29E+11 | 1,00E-01 |
|  | rRNA metabolic process (GO:0016072) | 4,83E+12 | 1,06E-01 |
|  | rRNA processing (GO:0006364) | 6,51E+11 | 1,06E-01 |
|  | regulation of RNA polymerase II transcription preinitiation complex assembly (GO:0045898) | 6,96E+11 | 1,06E-01 |
|  | regulation of phosphatidylinositol 3-kinase signaling (GO:0014066) | 8,65E+11 | 1,13E-01 |
|  | ribosome biogenesis (GO:0042254) | 1,04E-03 | 1,18E-01 |
|  | pteridine-containing compound metabolic process (GO:0042558) | 1,20E-03 | 1,21E-01 |
|  | response to UV-A (GO:0070141) | 1,39E-03 | 1,27E-01 |
|  | negative regulation of cell cycle G2/M phase transition (GO:1902750) | 1,54E-03 | 1,28E-01 |
|  | negative regulation of G2/M transition of mitotic cell cycle (GO:0010972) | 2,35E-03 | 1,61E-01 |
|  | positive regulation of cellular catabolic process (GO:0031331) | 2,47E-03 | 1,61E-01 |
|  | positive regulation of G protein-coupled receptor signaling pathway (GO:0045745) | 2,59E-03 | 1,61E-01 |
|  | positive regulation of transferase activity (GO:0051347) | 2,95E-03 | 1,61E-01 |
|  | NIK/NF-kappaB signaling (GO:0038061) | 3,26E-03 | 1,61E-01 |
|  | regulation of canonical Wnt signaling pathway (GO:0060828) | 3,46E-03 | 1,61E-01 |
|  | positive regulation of phosphorylation (GO:0042327) | 3,46E-03 | 1,61E-01 |
|  | positive regulation of transcription initiation from RNA polymerase II promoter (GO:0060261) | 3,79E-03 | 1,61E-01 |
|  | regulation of cellular response to heat (GO:1900034) | 3,92E-03 | 1,61E-01 |
|  | axon extension (GO:0048675) | 4,12E-03 | 1,61E-01 |
|  | regulation of monooxygenase activity (GO:0032768) | 4,12E-03 | 1,61E-01 |
|  | regulation of protein phosphorylation (GO:0001932) | 4,28E-03 | 1,61E-01 |
|  | regulation of G protein-coupled receptor signaling pathway (GO:0008277) | 4,35E-03 | 1,61E-01 |
|  | ribosomal large subunit assembly (GO:0000027) | 4,47E-03 | 1,61E-01 |
| GO terms, up-regulated in aged cardiac FB upon senolytics | regulation of feeding behavior (GO:0060259) | 6,58E+11 | 2,74E-01 |
|  | positive regulation of developmental growth (GO:0048639) | 1,06E-03 | 2,74E-01 |
|  | small GTPase mediated signal transduction (GO:0007264) | 1,07E-03 | 2,74E-01 |
|  | regulation of dephosphorylation (GO:0035303) | 1,32E-03 | 2,74E-01 |
|  | keratan sulfate metabolic process (GO:0042339) | 1,42E-03 | 2,74E-01 |
|  | extracellular structure organization (GO:0043062) | 1,59E-03 | 2,74E-01 |
|  | external encapsulating structure organization (GO:0045229) | 1,64E-03 | 2,74E-01 |
|  | respiratory system development (GO:0060541) | 1,78E-03 | 2,74E-01 |
|  | positive regulation of feeding behavior (GO:2000253) | 2,16E-03 | 2,74E-01 |
|  | collagen fibril organization (GO:0030199) | 2,18E-03 | 2,74E-01 |
|  | negative regulation of transcription, DNA-templated (GO:0045892) | 2,21E-03 | 2,74E-01 |
|  | regulation of axon extension (GO:0030516) | 2,42E-03 | 2,74E-01 |
|  | regulation of cellular catabolic process (GO:0031329) | 2,87E-03 | 2,74E-01 |
|  | epithelial tube branching involved in lung morphogenesis (GO:0060441) | 3,21E-03 | 2,74E-01 |
|  | cyclic purine nucleotide metabolic process (GO:0052652) | 3,60E-03 | 2,74E-01 |
|  | positive regulation of axonogenesis (GO:0050772) | 3,82E-03 | 2,74E-01 |
|  | cellular response to osmotic stress (GO:0071470) | 4,13E-03 | 2,74E-01 |
|  | regulation of membrane protein ectodomain proteolysis (GO:0051043) | 4,13E-03 | 2,74E-01 |
|  | positive regulation of axon extension (GO:0045773) | 4,13E-03 | 2,74E-01 |
|  | negative regulation of nucleic acid-templated transcription (GO:1903507) | 4,43E-03 | 2,74E-01 |
|  | regulation of aldosterone biosynthetic process (GO:0032347) | 4,45E-03 | 2,74E-01 |
|  | Cdc42 protein signal transduction (GO:0032488) | 4,45E-03 | 2,74E-01 |
|  | chemical homeostasis within a tissue (GO:0048875) | 4,45E-03 | 2,74E-01 |
|  | endothelial cell morphogenesis (GO:0001886) | 4,45E-03 | 2,74E-01 |
|  | glandular epithelial cell development (GO:0002068) | 4,45E-03 | 2,74E-01 |
| GO terms, down-regulated in aged cardiac FB upon senolytics | ribosome biogenesis (GO:0042254) | 1,11E+04 | 8,60E+05 |
|  | rRNA metabolic process (GO:0016072) | 1,26E+04 | 8,60E+05 |
|  | ncRNA processing (GO:0034470) | 2,25E+04 | 1,02E+07 |
|  | rRNA processing (GO:0006364) | 3,29E+04 | 1,12E+07 |
|  | nuclear-transcribed mRNA catabolic process, nonsense-mediated decay (GO:0000184) | 4,84E+04 | 1,32E+08 |
|  | cytoplasmic translation (GO:0002181) | 9,76E+04 | 2,22E+08 |
|  | cellular macromolecule biosynthetic process (GO:0034645) | 2,05E+05 | 4,00E+07 |
|  | SRP-dependent cotranslational protein targeting to membrane (GO:0006614) | 1,34E+07 | 2,28E+09 |
|  | cotranslational protein targeting to membrane (GO:0006613) | 2,06E+07 | 3,12E+09 |
|  | peptide biosynthetic process (GO:0043043) | 3,23E+06 | 4,40E+09 |
|  | protein targeting to ER (GO:0045047) | 5,07E+06 | 6,29E+08 |
|  | nuclear-transcribed mRNA catabolic process (GO:0000956) | 5,96E+06 | 6,78E+08 |
|  | gene expression (GO:0010467) | 4,61E+08 | 4,84E+10 |
|  | cellular protein metabolic process (GO:0044267) | 5,47E+08 | 5,33E+10 |
|  | translation (GO:0006412) | 6,18E+08 | 5,62E+10 |
|  | ribosomal large subunit assembly (GO:0000027) | 1,01E+09 | 8,62E+09 |
|  | ribosome assembly (GO:0042255) | 1,87E+10 | 1,50E+12 |
|  | ribosomal large subunit biogenesis (GO:0042273) | 4,10E+09 | 3,11E+10 |
|  | regulation of inflammatory response (GO:0050727) | 2,46E+10 | 1,77E-03 |
|  | positive regulation of inflammatory response (GO:0050729) | 5,40E+10 | 3,68E-03 |
|  | regulation of translation (GO:0006417) | 5,76E+10 | 3,74E-03 |
|  | ribonucleoprotein complex assembly (GO:0022618) | 7,24E+10 | 4,49E-03 |
|  | response to cytokine (GO:0034097) | 1,34E+12 | 7,93E-03 |
|  | positive regulation of cysteine-type endopeptidase activity involved in apoptotic process (GO:0043280) | 2,69E+12 | 1,53E-02 |
|  | activation of cysteine-type endopeptidase activity involved in apoptotic process (GO:0006919) | 3,49E+12 | 1,83E-02 |
| GO terms, up-regulated in aged cardiac PC upon senolytics | muscle contraction (GO:0006936) | 3,79E+06 | 9,56E+09 |
|  | regulation of release of sequestered calcium ion into cytosol by sarcoplasmic reticulum (GO:0010880) | 1,63E+08 | 2,06E+12 |
|  | regulation of heart contraction (GO:0008016) | 4,79E+08 | 4,03E+11 |
|  | heart contraction (GO:0060047) | 2,14E+10 | 1,35E-03 |
|  | regulation of cardiac muscle contraction by regulation of the release of sequestered calcium ion (GO:0010881) | 5,57E+10 | 2,34E-03 |
|  | membrane depolarization during cardiac muscle cell action potential (GO:0086012) | 5,57E+10 | 2,34E-03 |
|  | regulation of heart rate by cardiac conduction (GO:0086091) | 1,36E+11 | 4,90E-03 |
|  | cardiac muscle cell action potential (GO:0086001) | 1,94E+11 | 6,11E-03 |
|  | regulation of cardiac muscle contraction by calcium ion signaling (GO:0010882) | 2,21E+11 | 6,20E-03 |
|  | protein maturation (GO:0051604) | 2,47E+11 | 6,22E-03 |
|  | regulation of cardiac conduction (GO:1903779) | 2,71E+11 | 6,22E-03 |
|  | cardiac muscle contraction (GO:0060048) | 4,63E+10 | 9,18E-03 |
|  | canonical glycolysis (GO:0061621) | 5,09E+10 | 9,18E-03 |
|  | glucose catabolic process to pyruvate (GO:0061718) | 5,09E+10 | 9,18E-03 |
|  | glycolytic process through glucose-6-phosphate (GO:0061620) | 6,54E+10 | 1,10E-02 |
|  | striated muscle contraction (GO:0006941) | 1,57E+12 | 2,47E-02 |
|  | cell junction assembly (GO:0034329) | 1,87E+11 | 2,64E-02 |
|  | aerobic electron transport chain (GO:0019646) | 1,88E+12 | 2,64E-02 |
|  | mitochondrial ATP synthesis coupled electron transport (GO:0042775) | 2,10E+12 | 2,74E-02 |
|  | cardiac muscle cell proliferation (GO:0060038) | 2,35E+11 | 2,74E-02 |
|  | striated muscle cell proliferation (GO:0014855) | 2,35E+11 | 2,74E-02 |
|  | mitochondrial respiratory chain complex I assembly (GO:0032981) | 2,61E+11 | 2,74E-02 |
|  | muscle organ development (GO:0007517) | 2,61E+11 | 2,74E-02 |
|  | NADH dehydrogenase complex assembly (GO:0010257) | 2,61E+11 | 2,74E-02 |
|  | mitochondrial respiratory chain complex assembly (GO:0033108) | 2,85E+11 | 2,88E-02 |
| GO terms, down-regulated in aged cardiac PC upon senolytics | gene expression (GO:0010467) | 9,01E+09 | 1,15E-02 |
|  | nuclear-transcribed mRNA catabolic process, nonsense-mediated decay (GO:0000184) | 4,41E+10 | 2,03E-02 |
|  | ncRNA processing (GO:0034470) | 4,75E+09 | 2,03E-02 |
|  | peptide biosynthetic process (GO:0043043) | 6,36E+10 | 2,04E-02 |
|  | nuclear-transcribed mRNA catabolic process (GO:0000956) | 9,29E+10 | 2,13E-02 |
|  | SRP-dependent cotranslational protein targeting to membrane (GO:0006614) | 1,05E+11 | 2,13E-02 |
|  | cytoplasmic translation (GO:0002181) | 1,25E+12 | 2,13E-02 |
|  | cotranslational protein targeting to membrane (GO:0006613) | 1,33E+12 | 2,13E-02 |
|  | DNA replication-independent nucleosome organization (GO:0034724) | 1,51E+11 | 2,15E-02 |
|  | cellular protein metabolic process (GO:0044267) | 1,95E+12 | 2,50E-02 |
|  | protein targeting to ER (GO:0045047) | 2,20E+11 | 2,56E-02 |
|  | cellular macromolecule biosynthetic process (GO:0034645) | 3,03E+11 | 3,23E-02 |
|  | regulation of RNA splicing (GO:0043484) | 4,26E+11 | 3,92E-02 |
|  | translation (GO:0006412) | 4,28E+10 | 3,92E-02 |
|  | toll-like receptor 2 signaling pathway (GO:0034134) | 6,74E+10 | 5,75E-02 |
|  | ribosome assembly (GO:0042255) | 7,83E+11 | 6,26E-02 |
|  | regulation of mRNA splicing, via spliceosome (GO:0048024) | 9,23E+11 | 6,95E-02 |
|  | regulation of alternative mRNA splicing, via spliceosome (GO:0000381) | 1,05E-03 | 7,45E-02 |
|  | clathrin-dependent endocytosis (GO:0072583) | 1,27E-03 | 8,53E-02 |
|  | nucleosome assembly (GO:0006334) | 1,37E-03 | 8,53E-02 |
|  | amino sugar catabolic process (GO:0046348) | 1,40E-03 | 8,53E-02 |
|  | proteoglycan biosynthetic process (GO:0030166) | 2,13E-03 | 1,24E-01 |
|  | rRNA metabolic process (GO:0016072) | 2,35E-03 | 1,31E-01 |
|  | sterol metabolic process (GO:0016125) | 2,74E-03 | 1,35E-01 |
|  | ESCRT complex disassembly (GO:1904896) | 2,95E-03 | 1,35E-01 |

